# Leveraging TNFR2 for antitumour immunity: T reg depletion and myeloid reprogramming versus T cell costimulation

**DOI:** 10.64898/2026.01.04.697572

**Authors:** Linda Mårtensson, Petra Holmkvist, Kirstie Cleary, Carolin Svensson, Monika Semmrich, Niyaz Yoosuf, Alexandra Gabriela Ferreira, Mathilda Kovacek, Mimoza Bodén, Therese Blidberg, Osman Dadas, Jenny Mattsson, David Ermert, Elin Birgersson, Vincentiu Pitic, Martin Taylor, Mona Yazdani, Ulla-Carin Tornberg, Ingrid Karlsson, Sean Lim, Stephen A Beers, Mark Cragg, Björn Frendéus, Ingrid Teige

## Abstract

Although checkpoint inhibitors have revolutionized cancer treatment, responses are largely restricted to targeting CTLA-4 and PD-(L)1. TNFR2 has been identified as a target of interest, but a lack of understanding of whether agonists or blockers should be used, and whether FcγR interactions promote efficacy, has hampered therapeutic antibody development. Here, we engineered two distinct α-TNFR2 types: ligand-blocking FcγR-engaging versus non-ligand-blocking agonist antibodies. Both showed potent anti-tumor efficacy across multiple syngeneic mouse tumor models and combined effectively with α-PD-1. The ligand-blocking antibody depleted intratumoral T regs and reprogrammed the myeloid compartment, directing monocytes towards a pro-inflammatory macrophage and dendritic cell trajectory. Surprisingly, this effect depended on both inhibitory and activating FcγRs. In contrast, the agonist directly co-stimulated T and NK cells, through partially FcγR-independent mechanisms. Both antibodies converged on activating tumor-specific CD8^+^ T cells mediating tumor rejection. Clinical assessment of both ligand-blocking (BI-1808) and agonist (BI-1910) α-TNFR2 antibodies is currently ongoing.

**Graphical abstract:**

- Ligand-blocking anti-TNFR2 mAbs (BI-1808 and 3F10) deplete CCR8^+^ T regs and activate myeloid cells, remodeling the TME and leading to CD8^+^ T cell killing of cancer cells. T reg depletion and myeloid activation are mediated through interaction with activating FcγRs and the inhibitory FcγRIIB, respectively.
- Agonist anti-TNFR2 mAbs (BI-1910 and 5A05) intrinsically agonize TNFR2 on T cells and NK cells, which is enhanced by FcγRIIB crosslinking. The resulting broad lymphocyte activation mediates cancer cell killing.

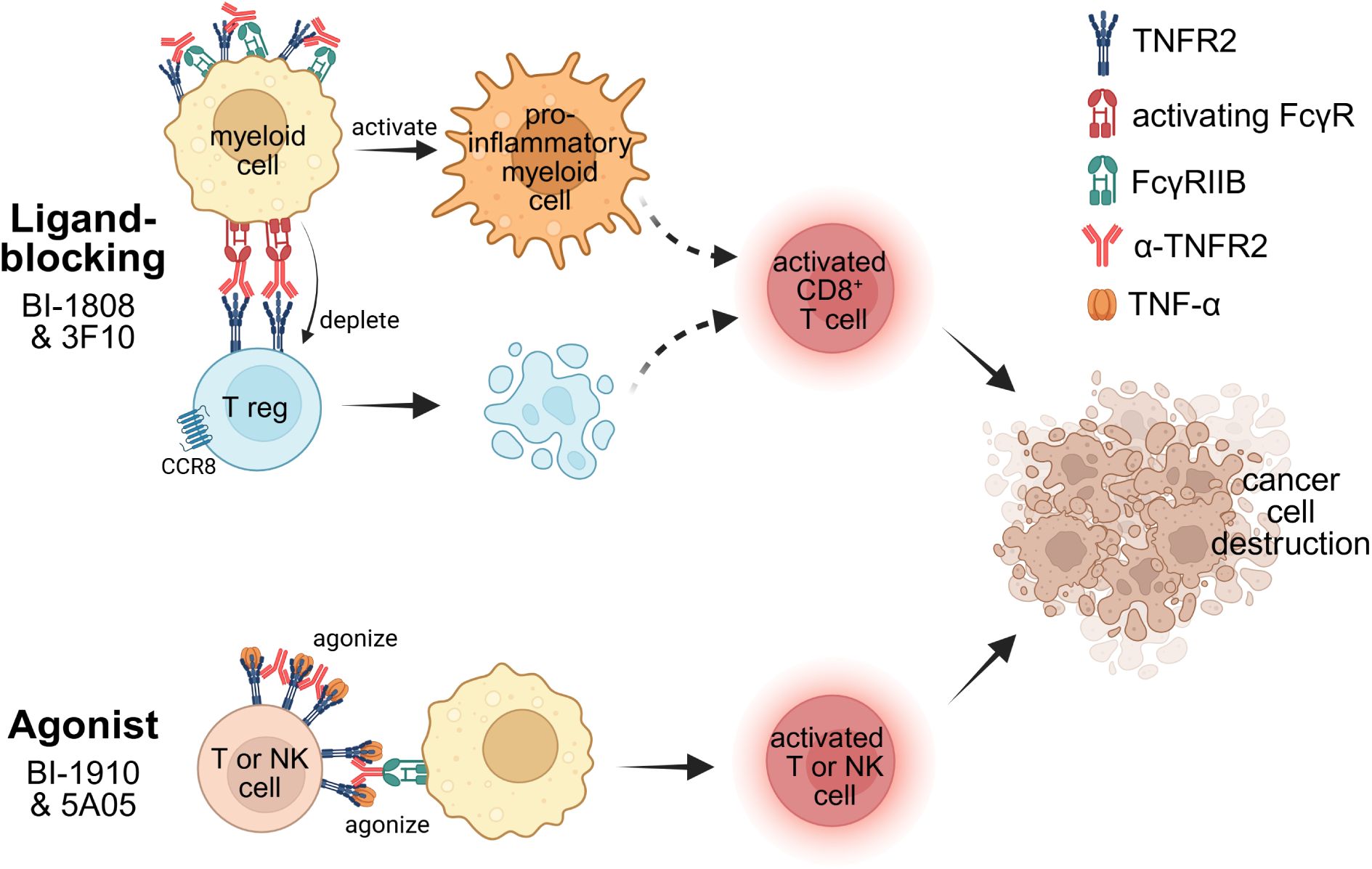

## Introduction

Immune checkpoint blockade (ICB) against PD-1, PD-L1, or CTLA-4 has transformed cancer care and is now used to treat over thirty different cancer types ^1,2^ with more being evaluated ^3,4^. Still, many patients fail to respond, relapse and/or acquire resistance ^5^. Accordingly, there is a need to identify effective new therapeutic targets and mechanisms that combine with existing ICBs to induce robust anti-tumor immunity and improve clinical responses in more patients. Despite significant efforts in the field, only a single antibody (relatlimab) targeting a new immune checkpoint (LAG-3) has been approved by the FDA in the last 10 years. Of note, relatlimab is approved only for use with α-PD-1 without evidence of single-agent activity ^6^.

One immunomodulatory target receiving increasing attention is TNFR2 ^7–9^. TNFR2 shares ligands (TNF-α and LT-α) with the highly related receptor TNFR1, with both receptors resulting in NF-κB activation and immune modulation ^10^. TNFR2 expression is largely restricted to immune cells. Its role is complex, with both pro- and anti-inflammatory activities depending on context. As a result, TNFR2 agonists have been proposed as treatments for both chronic inflammation and cancer ^8,11–14^. Gene deletion studies show that a lack of TNFR2 increases incidence and severity in inflammatory models ^15,16^, suggesting that TNFR2 largely mediates anti-inflammatory signals. High expression of TNFR2 on immunoregulatory cells such as T regs and myeloid derived suppressor cells, also supports that TNFR2 mainly works to inhibit inflammation, and several publications demonstrate an important role of TNFR2 in T reg proliferation and survival ^14,17–21^. Accordingly, the first TNFR2-targeting agents were explored for regulating autoimmunity and inflammation through T reg activation ^22,23^, with cancer therapeutics proposed as blocking antibodies either by abrogating T reg functions or through T reg deletions ^24–27^. However, contrasting data demonstrate that TNFR2 acts as a costimulatory factor also for conventional T cells ^28–30^. In fact, all effector T cells display high TNFR2 expression ^31,32^. Moreover, current evidence suggests that TNFR2 acts as a costimulatory receptor on multiple immune cells, much like other members of the TNFR superfamily such as 4-1BB, OX40, and CD27, whereby the net immune effect depends on factors such as individual cell expression, activation status, and immune cell subpopulation composition ^33^. Consequently, several recent publications have proposed TNFR2 agonists for cancer treatment ^7,8,34^.

Given the complexity of TNF biology, we wished to further investigate the anti-cancer potential of TNFR2 antibodies and define the key characteristics required for optimal efficacy. Accordingly, we generated two panels of antibodies targeting mouse or human TNFR2 for exploration in immunocompetent in vivo models and therapeutic development, respectively. Furthermore, we identified and characterized two lead antibodies in each panel – one ligand-blocking and one agonistic non-blocking antibody. Importantly, as part of these studies, we evaluated antibody isotype and FcγR-dependency, known to influence antibodies’ mode of action through differential interaction with activating or inhibitory FcγRs ^35,36^. Once in their optimal isotype, these antibodies were evaluated for anti-tumor efficacy and mode of action in vivo, in comparison to, and in combination with, mAbs directed to existing ICB, CTLA-4 and PD-1.

We show that TNFR2 ligand-blocking antibodies simultaneously mediate FcγR-dependent deletion of tumor-infiltrating T regs and FcγR-dependent agonism of myeloid cells, and that both mechanisms are essential for strong anti-tumor activity. In contrast, non-ligand-blocking agonists activate T cells and NK cells, resulting in tumor regressions. These studies shed light on the previous uncertainties around TNFR2 as a target for cancer and demonstrate how both ligand-blocking, as well as agonist α-TNFR2 antibodies, can be optimally engineered for clinical translation.

## Results

### Generation and in vitro characterization of human and murine α-TNFR2 antibodies

To identify potentially therapeutic TNFR2 antibodies, we generated a wide panel of human TNFR2-targeting antibodies using our proprietary phage-display library nCoDeR ^37^. The antibodies spanned fully ligand-blocking (e.g. BI-1808) to non-blocking clones (e.g. BI-1910; Fig. 1a-b). These clones were further characterized in a series of additional assays (data not shown), and based on these data, BI-1808 and BI-1910 were chosen for further investigation. Both antibodies showed potent binding to TNFR2-expressing activated primary T cells (Fig. 1c) and TNFR2 protein, but not to the closest TNFR2 homologs LTβR and TNFR1 (Extended Data fig.1a-c).

**Figure 1:**
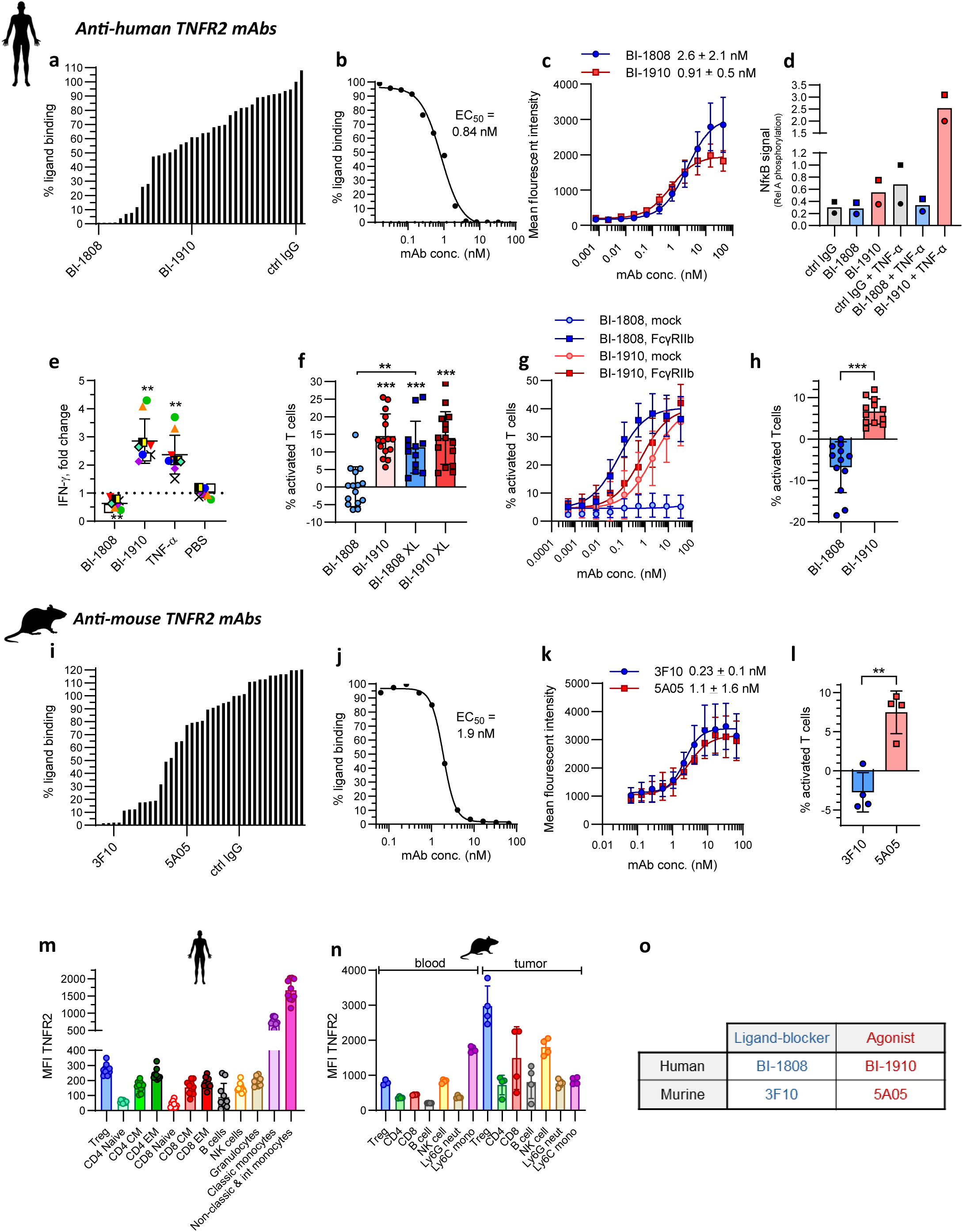
Generation and in vitro characterization of human and murine anti-TNFR2 antibodies a, Ligand-blocking capacity of generated mAbs targeting human TNFR2, BI-1808 and BI-1910 are indicated. **b,** 5 nM TNF-a binding to TNFR2 in presence of increasing amounts of BI-1808 **c,** Binding properties of the two lead clones to in vitro activated CD4^+^ T cells. Representative curves from 3 donors are shown. The EC_50_ value is the mean ± SD of T cell binding to the totally 26 and 22 donors assayed respectively. **d,** Western blot data showing canonical NF-kB signaling in TNFR1^-^/^-^ NK cells as studied by RelA phosphorylation. The figure shows two independent experiments; individual values are indicated either by circles or squares. **e,** NK cell activation as measured by IFN-g release. **f,** CD4^+^ T cell activation as measured by proliferation in the presence or absence of anti-human Fab cross-linking (XL). **g,** TNFR2 transfected NF-kB reporter Jurkat T cells, co-cultured with CHO cells lacking (mock) or expressing FcgRIIB (FcgRIIB), showing NF-kB activation as measured by GFP positivity. **h,** Co-culture of CD3/CD28 bead stimulated T cells and tumor associated macrophage-like cells. T cell activation measured by proliferation and CD25 upregulation. For figure e, f and h, figures show mean values of at least three independent experiments ± SD and each dot is one donor. Since baseline values vary between individual donors, each donor is normalized to the mean of ctrl IgG. **= p<0,01 ***= p<0,001 using one-way ANOVA. **i,** Ligand-blocking capacity of corresponding mAbs targeting murine TNFR2. 3F10 and 5A05 surrogates are indicated. **j,** 2 nM TNF-a binding to mTNFR2 in presence of increasing amounts of 3F10. **k,** Binding properties of the two chosen surrogate clones on in vitro activated CD4^+^ T cells. The figure shows representative binding curves from 2 independent experiments. The EC_50_ ± SD shows values of binding to T cells from 11 and 5 donors, respectively. **l,** Co-culture of tumor-associated macrophage-like cells and T cells. T cell activation measured by proliferation. Since baseline varies between individual donors, each donor is normalized to mean of the IgG ctrl. Figures show mean values ± SD, from two independent experiments. **= p<0,01 as calculated using one-way ANOVA. **m,** TNFR2 expression on human immune cells from fresh whole blood. **n,** TNFR2 expression on murine immune cells from fresh whole blood or cells isolated from CT26 tumors. **o,** Summary of human lead candidate mAbs and their murine surrogates.

To study their effect on TNF-α:TNFR2-signaling, we used a TNFR2-expressing NK cell line lacking TNFR1. In these cells, the ligand-blocking antibody BI-1808 blocked TNF-α-induced NF-κB activation and did not itself induce NF-κB. BI-1910, in contrast, induced NF-κB, with activity further increased in the presence of TNF-α (Fig. 1d). Therefore, BI-1910 acted as an agonist and BI-1808 as an antagonist. When combined, they neutralized each other (Fig. S1d).

We next studied the effects of the antibodies on TNFR2-expressing immune cells. First, we assessed the ability of the mAbs to augment IFN-γ production from NK cells ^38^. Again, BI-1808 behaved as an antagonist, lowering the IFN-γ levels, and BI-1910 functioned as an agonist, increasing IFN-γ (Fig. 1e). The antibodies had similarly opposed effects on T cell proliferation. BI-1910 acted as a strong agonist, whereas BI-1808 blocked proliferation (Fig. 1f-g). To mimic the in vivo situation where FcγR (in particular FcγRIIB) can elicit mAb cross-linking ^36,39,40^ we performed assays with anti-antibodies or FcγRIIB-expressing CHO cells. While the agonistic effect of BI-1910 was unaffected, BI-1808 was converted from an antagonist to an agonist by both approaches (Fig. 1f-g). Finally, using CD3/CD28 stimulated T cells and FcγR-expressing tumor-associated macrophage-like cells (Fig. 1h), BI-1910 but not BI-1808 provided TNFR2-mediated co-stimulatory activity (Fig. 1h). Together, these data demonstrated that in vitro BI-1910 acted as an intrinsic agonist (independent of FcγR or other cross-linking) while BI-1808 acted as a ligand-blocking antagonist with the potential for FcγR-dependent co-stimulatory activity under sufficient cross-linking.

To enable the translation of these data to the immune-competent in vivo tumor setting, we generated an equivalent panel of murine TNFR2-targeting antibodies with the same characteristics, ranging from full ligand blockers to non-blockers (Fig. 1i-j). These antibodies were tested in assays similar to those described above, enabling us to identify murine surrogates that closely parallelled the human lead candidates: a ligand-blocking antibody (3F10) and a non-blocking agonist (5A05). They displayed similar high specificity for TNFR2, binding properties, and analogous functional activity to their human counterparts (Fig. 1i-l and Extended Data fig.1e-g). Importantly, human and murine immune cells showed very similar TNFR2 expression, see Fig. 1m-n, supporting the use of mice for in vivo mode-of action experiments. A summary of the investigated mAbs is shown in Fig. 1o.

### Ligand-blocking and agonistic antibodies have broad anti-tumor activity acting through divergent isotypes and FcγR

Next, we tested the anti-tumor efficacy of the ligand-blocking antibody (3F10) and a non-blocking agonist (5A05) in immunocompetent mice transplanted with CT26 tumors, concurrently evaluating antibody isotype, which is known to influence the in vivo mode of action through differential interaction with activating or inhibitory FcγRs ^35,36,41,42^ . We therefore engineered mAb with Fc mutations known to impair FcγR interaction (mIgG1 N297A, here referred to as Fc null) and animals expressing activating, inhibitory, both, or no FcγRs.

Perhaps surprisingly, both the ligand-blocking and agonistic antibodies conferred strong anti-tumor activity, as demonstrated by tumor growth inhibition, regression, and prolonged animal survival (Fig. 2a-d). 3F10 showed robust anti-tumor activity in antibody formats known to be optimal for either agonizing TNFRs (mIgG1, low A:I ratio) ^43^ or target cell depletion (mIgG2a, high A:I ratio) (Fig. 2a-d). In contrast, the agonist 5A05 was clearly most effective in the mIgG1 format. Whilst the ligand-blocking antibodies’ (3F10) therapeutic effect was lost in the Fc null format, the agonist (5A05) retained significant anti-tumor activity (Fig. 2a-d). The ligand-blocking antibodys’ unique requirement for activating and/or inhibitory FcγR for efficacy was confirmed in CT26 tumor-bearing mice lacking either the inhibitory FcγRII (FcγRII^-^/^-^), all activating FcγRs (γ-chain^-^/^-^), or both (γ-chain^-^/^-^ x FcγRII^-^/^-^) (Fig. 2e-f). Collectively, these observations indicated that the ligand-blocking antibody engaged both activating and inhibitory FcγRs to deliver optimal therapeutic effects, with the high A:I ratio mIgG2a isotype being most efficacious. In contrast, the agonistic antibody had intrinsic activity that was enhanced and most effective in the FcγRII-engaging low A:I ratio mIgG1 format. Therefore, unless stated otherwise, we hereafter used these optimal mAb formats; mIgG2a for 3F10 and mIgG1 for 5A05.

**Figure 2:**
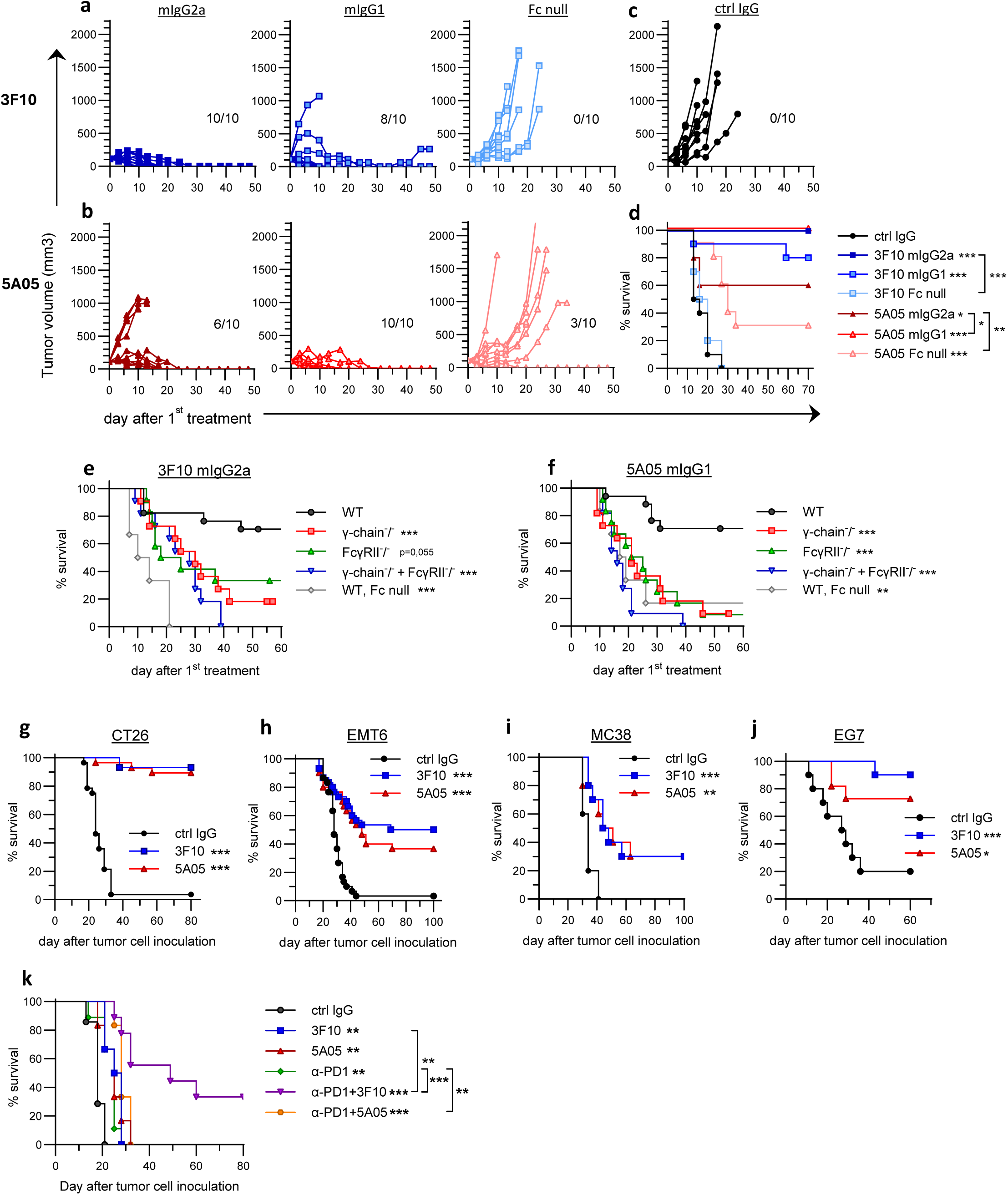
Both ligand-blocking and agonistic antibodies have broad anti-tumor activity but require different antibody isotypes for optimal efficacy. **a,** Tumor growth and survival in CT26 tumor-bearing mice treated with the ligand-blocking antibody 3F10 or **b,** the non-ligand-blocking agonist 5A05 in different isotypes. **c,** Tumor growth in control IgG treated tumors and **d,** survival in all groups. Numbers in the graph indicate totally cured / total number of mice in the group. **e-f)** Survival in CT26 tumor bearing mice of various FcgR^-/-^ backgrounds treated with 3F10 or 5A05 respectively. Tumors were ca 100 mm^2^ at start of treatment. Survival of tumor-bearing mice treated with 3F10 or 5A05 showing **g,** combined data from three experiments (N=28-29 mice/group), tumors were 100-140 mm^3^ at the start of treatment. **h,** Combined data from three experiments (N=30 mice/group), tumors were 80-150 mm^3^ at the start of treatment. **i,** Data from one representative experiment (N=10 mice/group), tumors were 40-70 mm^3^ at the start of treatment. **j,** Data from two different experiments (N=10 mice/group), tumors were 100-150 mm^3^ at the start of treatment. **k,** Data from one representative experiment (N=6-9 mice/group) when tumors were ca. 30 mm^3^ at the start of treatment. For statistics in d-k) *= p<0,05 **= p<0,01 ***= p<0,001 using Log-rank (Mantel-Cox) test. Comparisons were made against control IgG unless otherwise indicated.

We next assessed activity in tumor models differing in CD8^+^ T cell infiltration and responsiveness to PD-1/ CTLA-4 ICB. Both antibodies had single-agent activity (Fig. 2g-j), which correlated with T cell infiltration (CT26 ≥ MC38 > EMT6 >>B16 ^44^, see Extended Data fig.2a-b), indicating immunomodulatory mechanisms of action. This notion was further supported by similar in vivo anti-tumor activities against TNFR2-expressing and TNFR2-/- syngeneic tumors (Extended Data fig.3a-b).

Significant anti-tumor activity was also observed in the T cell lymphoma model EG7. Due to the broad use of α-PD-1 in clinical practice, it is important to understand the combinatorial effects with PD-1 blockade. As shown in Extended Data fig.4a-b, both α-TNFR2 antibodies showed excellent combinatorial activity with α-PD-1. Uniquely, the ligand-blocking 3F10 antibody induced enhanced survival of animals in the poorly CD8^+^ T cell-infiltrated B16 model, which is resistant to α-PD-1 and α-CTLA-4 (Fig 2k).

### Combining ligand-blocking and agonistic TNFR2 antibodies compromises anti-tumor activity

Our data demonstrated that the two α-TNFR2 antibody types showed similar anti-tumor activity in several experimental models but differed in in vitro and in vivo functional characteristics. This indicated that they triggered anti-tumor immunity by, at least partially, different mechanisms. To assess whether these mechanisms enhanced or compromised each other, we evaluated 3F10 and 5A05 in combination. When combined, tumor control and animal survival were reduced compared to single-agent treatments, consistent with divergent, potentially competing, mechanisms of action (Fig. 3a-b). Importantly, both antibodies bound TNFR2 independently. Competitive target binding could therefore not account for the diminished combinatorial anti-tumor activity (Extended Data fig.5a-b).

**Figure 3.**
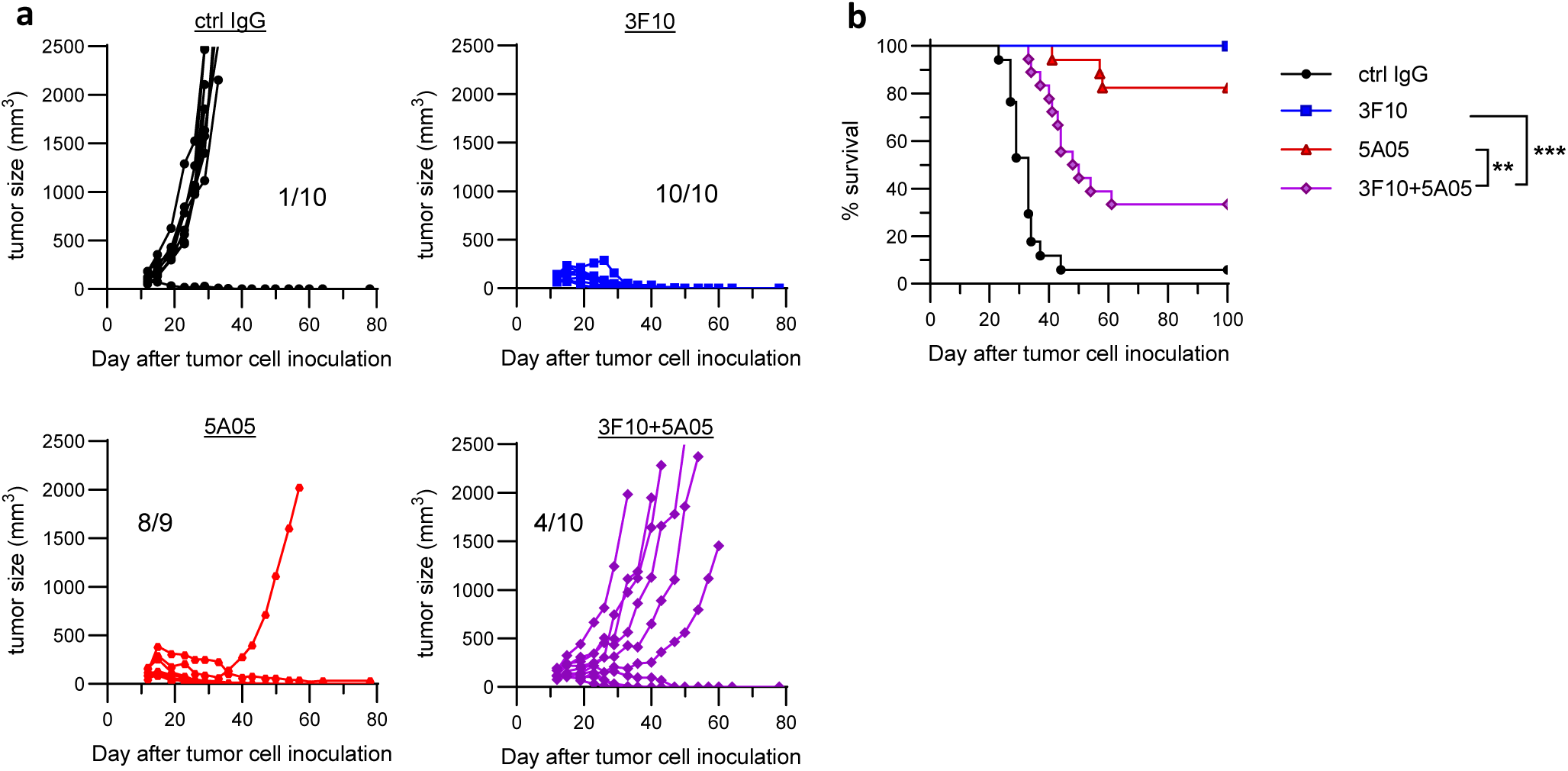
Combining ligand-blocking 3F10 and agonist 5A05 nullifies each other’s anti-tumor effect a, Tumor growth curves from CT26 tumor-bearing mice treated with 3F10, 5A05 or a combination of both, data show one representative experiment. Numbers in the graph indicate totally cured / total number of mice in the group. **b,** Survival curve from two independent experiments combined, 17-18 mice / group. The tumors were ca 100-150 mm^3^ at the start of treatment.

### α-TNFR2 antibodies elicit divergent transcriptional programs in tumor-infiltrating T cells

Next, we performed single-cell RNA sequencing (scRNA) to agnostically characterize the immunomodulatory effects of the two antibodies in the CT26 model, comparing them with α-CTLA-4 and α-PD-1 treatment. CD45^+^ cells were isolated from tumors on day 2 and 8 after the first treatment to capture early events and those coinciding with tumor regression, respectively (Fig. 4a). The integrated analyses across all groups identified several immune cell populations (Extended Data fig.6a).

**Figure 4:**
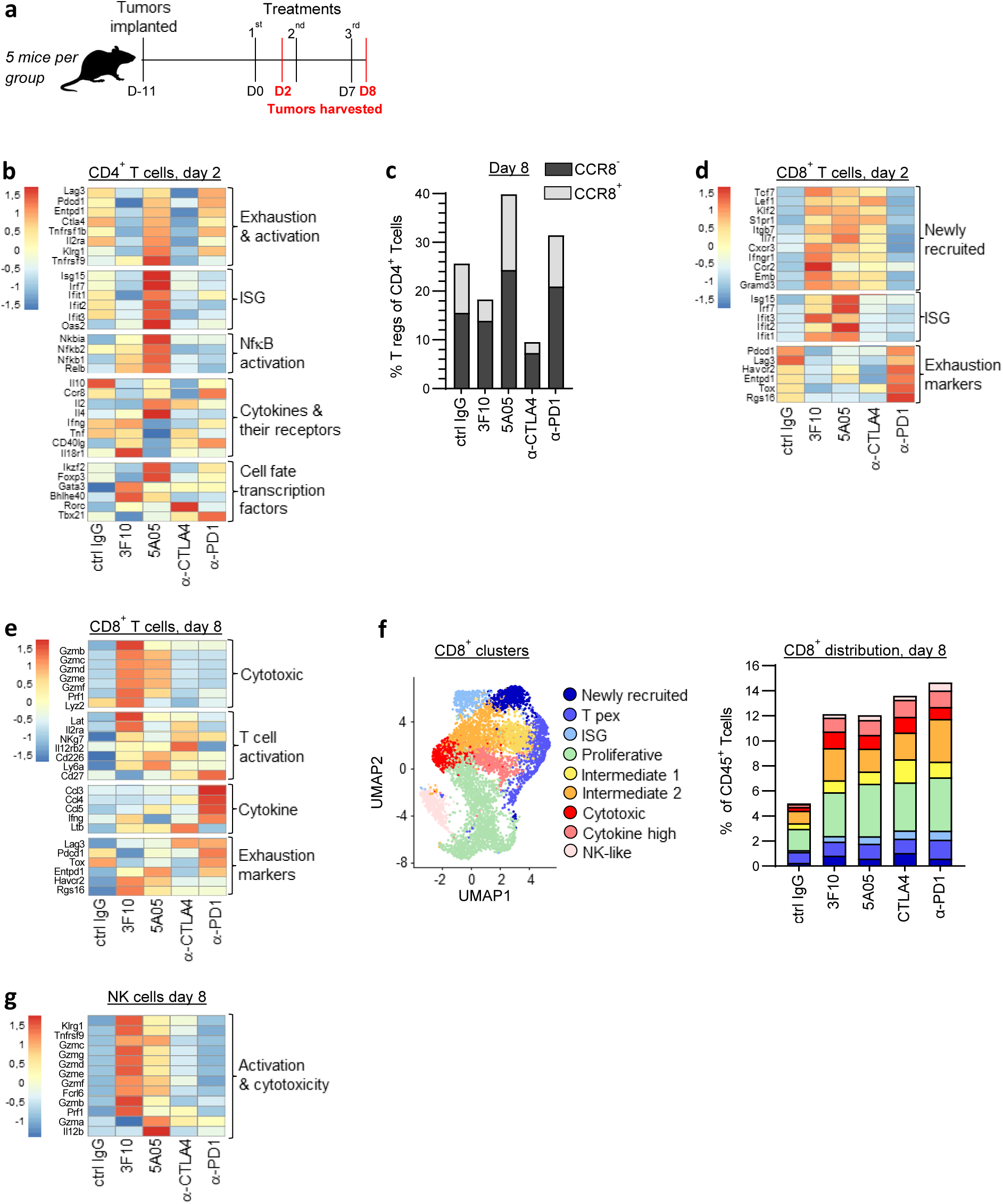
a-TNFR2 antibodies elicit divergent transcriptional programs in CD4^+^ T cells and activate cytotoxic CD8^+^ T cells a, Schematic layout of the experimental setup. **b-g,** Single-cell RNA sequencing data from CD45^+^ sorted cells isolated from five CT26 tumors per treatment group. Tumors were harvested on day 2 and day 8 after first treatment as indicated. **b,** Heatmap on CD4^+^ T cell showing relative average expression of genes associated with activation, ISG signature, cytokines and transcription factors associated with cell fates e.g. T reg, Th2 and Th1. **c,** Percentage of *Ccr8*^-^ and *Ccr8*^+^ T regs of the CD4^+^ population. **d,** Heatmaps on CD8^+^ T cells showing relative average expression of genes associated with newly recruited T cells, ISG signature, and exhaustion markers for different treatments at day 2 or **e,** genes associated with cytotoxicity, T cell activation, cytokines and exhaustion for different treatments at day 8. **f,** UMAP plot of all CD8^+^ T cell clusters and the bar-plot showing distribution as percentage of total CD45^+^ cells on day 8. **g,** Heatmap on NK cell showing relative average expression of genes associated with activation and cytotoxicity.

Several key findings relating to lymphocytes were made. Firstly, the agonist 5A05 uniquely activated the CD4^+^ T cell population, with effects being most pronounced on day 2 (Fig. 4b). The induction of interferon-stimulated genes (ISG) and NfκB activation was particularly strong. The agonist preferentially induced Th2 cytokines with high IL-4 and low IFN-γ. Secondly, while 5A05 increased expression of T reg-associated genes such as Foxp3 and Ikzf2 (Helios), 3F10 reduced these, showing a clear dichotomy between the two TNFR2 antibodies. Quantifying T reg cells based on transcription pattern confirmed this (Fig. 4c). Interestingly, the Foxp3^+^ T reg population was split into two clusters, best divided by differential Ccr8 expression (Extended Data fig.6b-c). The *Ccr8*^+^ population was characterized by high expression of several activation and exhaustion markers (Extended Data fig.6d). Interestingly, 3F10 depleted only the *Ccr8*^+^ T regs while α-CTLA-4 reduced both populations (Fig. 4c).

Thirdly, 3F10, 5A05, and the α-CTLA-4 mAb all induced genes associated with the influx of newly recruited and proliferating stem-like CD8^+^ T cells on day 2, downregulating genes associated with exhaustion (Fig. 4d). In addition, both TNFR2 targeting antibodies increased ISG expression. By day 8, during tumor regression, the TNFR2 antibodies, especially 3F10, robustly induced a cytotoxic gene expression profile. In contrast, α-PD1 induced expression of cytokine genes (Fig. 4e). Quantifying cells in each cluster, all therapeutic antibodies had a similar overall increase in CD8^+^ T cells with a 4-5-fold increase in the cytotoxic and cytokine producing effector cells compared to isotype control (Fig. 4f). For genes used to classify CD8^+^ T cell clusters see Extended Data fig.6e. Finally, only the α-TNFR2 antibodies induced strong NK cell activation, suggesting that NK cells might uniquely contribute to their mode of action (Fig. 4g).

In summary, the agonist 5A05 specifically activated CD4^+^ T cells, including T regs, while the ligand-blocker 3F10 deleted T regs and activated a strong cytotoxic program in CD8^+^ T cells. Both antibodies increased CD8^+^ T cell recruitment and NK cell activation.

### Immune modulation by 3F10 is FcγR-dependent and results in tumor-specific CD8^+^ T cell responses

To confirm the above findings and further understand the role of FcγRs, we repeated the setup outlined in Fig. 4a, analyzing immune cells from tumors, draining lymph nodes, and spleens of CT26-bearing mice using flow cytometry (FCM). These data confirmed that 3F10 induced specific depletion of intratumoral CCR8^+^ T regs, while 5A05 increased the same T reg population (Fig. 5a). Both antibodies subsequently converged in their CD8^+^ T cell response; dramatically increasing CD8^+^ T cells overall (Fig. 5b), resulting in a 3 and 7-fold improved CD8^+^ T cell/T reg ratio (Fig. 5c). We also observed increases in tumor-specific CD8^+^ T cells (Fig. 5d) and a pronounced cytotoxic response seen with 3F10 as measured by Granzyme B^+^ CD8^+^ T cells (Fig. 5e). Upon tumor antigen stimulation ex vivo, α-TNFR2 treatment resulted in increased effector cytokine (IFN-γ, TNF-α) producing CD8^+^ T cells (Fig. 5f). Looking at T cell proliferation in tumor draining lymph nodes, it was again clear that both antibodies increased CD8^+^ T cell expansion while the agonist 5A05 had a greater effect on CD4^+^ T cells (Fig. 5g). These effects were largely FcγR-dependent since solely blocking TNFR2 using the Fc null variant of 3F10 had minimal impact.

**Figure 5:**
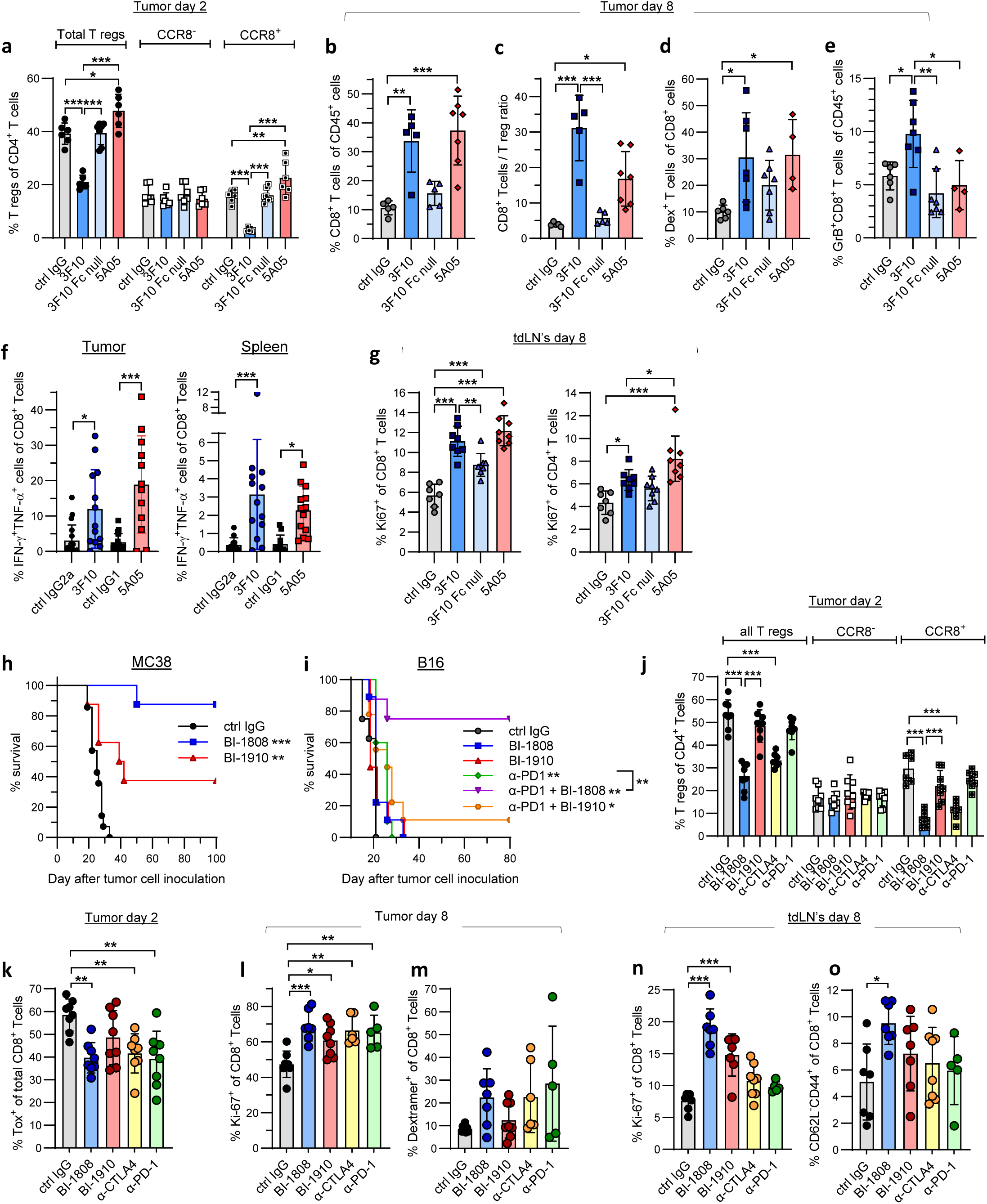
Immune cell modulations are FcgR dependent and result in tumor-specific CD8 ^+^ T cell responses. a-g) Tumors and draining lymph nodes were dissected from CT26 bearing mice on day 2 and 8 after the first treatment as indicated and analyzed by FCM. Each dot represents one individual mouse. **a,** Percentage of all or CCR8^-^ versus CCR8^+^ intratumoral T regs of the CD4^+^ T cell population. **b,** Results from day 8 showing the fraction of CD8^+^ T cells of infiltrating immune cells **c,** the CD8^+^ T cell to T reg ratio **d,** the percentage of tumor-specific CD8^+^ T cells, as analyzed by AH1 dextramer positivity and **e,** percent of Granzyme B positive CD8^+^ T cells of infiltrating immune cells. **f,** Antigen-specific effector recall T cell responses on day 10 after the start of treatment. Single cell suspensions from tumor or spleen were stimulated with AH1 peptide and analyzed for intracellular TNF-a and IFN-g. Data show the mean ± SD from 3 independent experiments. **g,** Fraction of proliferating CD8^+^ respectively CD4^+^ T cells in tumor-draining lymph nodes on day 8. **a-g,** Show mean values ± SD. **h-i,** Survival curves of **h,** MC38 or **i,** B16 inoculated mice transgenic for human TNFR2 and treated as indicated. **j-o,** Tumors and draining lymph nodes were dissected from MC38-bearing hTNFR2 transgenic mice at days 2 and 8 after the first treatment as indicated and analyzed by FCM. Each dot represents one individual mouse. **j,** Percentage of total and CCR8^-^ and CCR8^+^ intra-tumoral T regs of the CD4^+^ population. **k,** Percentage of exhausted cells in the CD8^+^ T cell population as measured by TOX, day 2. **l,** Percentage of proliferating CD8^+^ T cells and **m,** tumor-specific CD8^+^ T cells as measured by specific dextramers on day 8. **n-o,** Proliferative CD8^+^ T cells and effector memory cells in tumor-draining lymph nodes on day 8. j-o, Show mean values ± SD. For statistics in b-g, and k-o, one-way ANOVA; and for a, and j, two-way ANOVA were used. *=p<0,05, **=p<0,01 and ***=p<0,001 using Tukey’s adjustment for multiple comparisons. In h, tumors were ca 6×6 mm and in I, ca 4×4 mm at start of treatment, *= p<0,05 **= p<0,01 using Log-rank (Mantel-Cox) test. Comparisons were made against control IgG unless otherwise indicated.

The instrumental role of the activated CD8^+^ T cells was demonstrated in experiments where CD8 depletion completely abrogated the anti-tumor effect of both TNFR2 mAbs (Extended Data fig.7a). Interestingly, although both 3F10 and 5A05 induced strong NK cell activation (Fig. 4g), only the agonist 5A05 showed partial NK cell dependence for efficacy (Extended Data fig.7b). NK and CD8 depletion were verified using FCM (Extended Data fig.7c-d).

### Human lead candidate and mouse surrogate α-TNFR2 antibodies show similar anti-tumor and immune modulatory effects

To confirm that the results obtained using the murine surrogates were valid for the human candidates (Fig. 1), we evaluated our lead hTNFR2 mAbs in hTNFR2 knock-in mice. Importantly, murine TNF-α can bind and signal through hTNFR2, leaving the TNFR:TNF axis intact^45^. Using these mice, we first established that we had excellent anti-tumor effects with the human lead candidates BI-1808 (ligand-blocker, equivalent of 3F10) and BI-1910 (agonist, similar to 5A05) (Fig. 5h-i), although here BI-1808 outperformed BI-1910. Notably, and in keeping with the mouse surrogate data, in the difficult-to-treat B16 model, BI-1808 in combination with α-PD-1 resulted in 75% cures, whilst α-PD-1 alone mediated only modest tumor growth delay (Fig. 5i). Next, we performed an analogous immune profiling experiment in MC38 tumors using α-CTLA-4 and α-PD-1 as comparators. The data mirrored the previous results using the surrogate mAbs 3F10 and 5A05 and showed that BI-1808 and α-CTLA-4 specifically depleted CCR8^+^ T regs (Fig. 5j). We also verified overall decreased exhaustion in CD8^+^ T cells as measured by fewer TOX^+^ cells in the treated groups (Fig. 5k). On day 8, there was an influx of proliferating CD8^+^ T cells alongside increased tumor-specific CD8^+^ T cells as measured by antigen-specific dextramers in all treatment groups (Fig. 5l-m). In tumor-draining lymph nodes, only the α-TNFR2 antibodies induced CD8^+^ T cell proliferation, and BI-1808 exclusively increased mature effector memory T cells (CD62L^-^ CD44^+^) (Fig. 5n-o).

### Ligand-blocking α-TNFR2 elicits myeloid re-programming towards a moDC and DC3 phenotype important for anti-tumor activity

Having established the two antibodies’ differential effects on TILs, we next assessed modulating activity on tumor-associated myeloid cells using the CT26 tumor model. Based on well-described gene expression patterns ^46–49^, we defined 11 myeloid populations. These included monocytes, interferon-stimulated macrophages (ISG), MoDCs, and DC3s ^46–49^. Furthermore, we identified two MHCII^high^ populations: one Ly6i^+^expressing Itgax/CD11c (Ly6i^+^ MHCII^high^) and one high in anti-inflammatory “M2” markers such as C1q, Mrc1, and FolR2 (C1q^+^ MHCII^high^). Three additional populations were all relatively high in similar “M2” markers but largely lacked MHCII (defined as Arg1^+^, C1q1^+^ and C1q1^+^ proliferating). Finally, one population was high in Nos2 and Mmps (Nos2^+^), and another intermediate population shared characteristics with several of the others (intermediate), see Fig. 6a and Extended Data fig.7f-g. Based on these transcriptional signatures, the myeloid cells were ordered in a spectrum where the DCs were considered as having the highest anti-tumor capacity, with MHCII^low^ cells high in “M2” markers classified as the strongest tumor-promoting (Extended Data fig.7f). Quantifying the myeloid populations revealed a striking reduction in the populations bearing “M2” markers in the 3F10 treated group, alongside a simultaneous increase in DCs and DC-like cells (Fig. 6a-b). A heat map of relative gene changes clearly shows 3F10 reduced M2 macrophage associated genes (Fig. 6c). Calculating a ratio between populations at the two ends of the spectrum - the two populations with the assumed most tumor promoting characteristics, and the three DC or DC-like populations - illustrated that 3F10 reversed their relative proportions, with a 6-fold change compared to control IgG (Fig. 6d). A trajectory analysis starting from monocytes showed four trajectories ending either in DCs, Nos2^+^, or two populations with “M2” markers – the C1q^+^ MHCII^high^ and the C1q^+^ proliferating (Fig. 6e). Predictably, the dominating trajectory in control-treated mice was towards the pro-tumorigenic M2-like macrophages. Strikingly, 3F10 directed myeloid cells differentiation towards the trajectory ending in either DCs or the Nos2^+^ population, fundamentally remodeling the TME from a tumor-supportive state to a tumor-rejecting phenotype. This was not observed with the 5A05 antibody, again illustrating differences in mode-of-action between the two TNFR2 targeting antibodies. A similar but less pronounced remodeling was induced by α-CTLA-4, in accordance with previous work ^48^.

**Figure 6:**
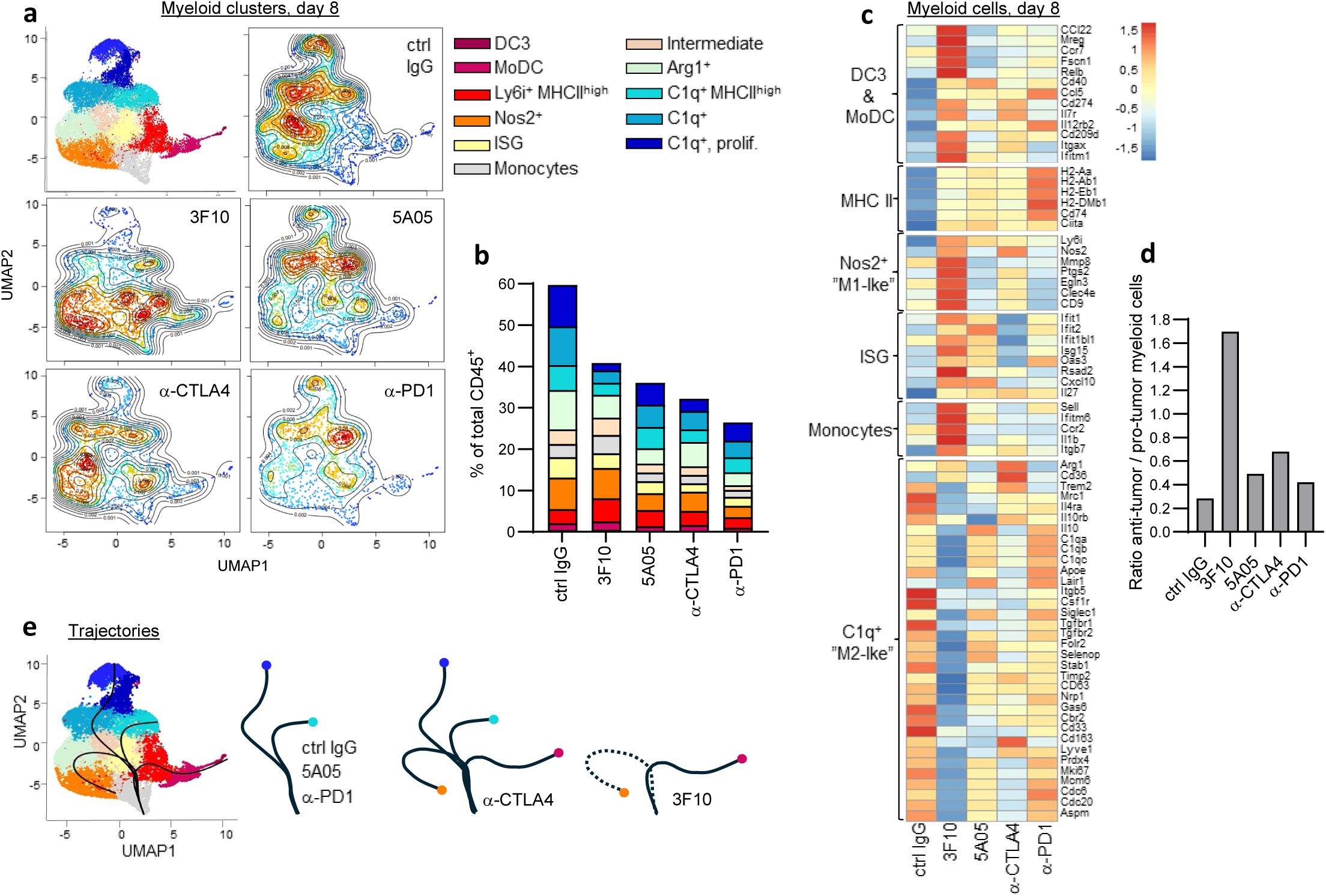
a-TNFR2 elicits myeloid programming towards a DC phenotype. a-e) Single-cell RNA sequencing data from CD45^+^ sorted cells isolated from five CT26 tumors per treatment group. The data shown is from tumors harvested on day 8 after the first treatment. **a,** UMAP plot of myeloid clusters colored by cell density for different treatment groups. **b,** The distribution of myeloid subpopulations as the percentage of total CD45^+^ cells. Pro-tumoral M2-like clusters are indicated in blue, and anti-tumoral M1-like and DC clusters are presented in red. **c,** Heatmap of relative average expression of selected myeloid genes for each treatment group. **d,** The ratio (calculated as the percentage of total CD45^+^ cells) between the two DC and Ly6i MHCII^high^ populations divided by the two C1q MHCII^low^ populations. **e,** Trajectory analysis of myeloid cells using slingshot shows four lineages starting in the monocytes.

To verify these findings, tumors from treated mice were harvested on D8 and analyzed by FCM. Seven myeloid or DC cell types were identified using established markers (Fig. 7a). A moDC population, likely also containing DC3 cells, was increased by both α-TNFR2 antibodies (Fig. 7b). However, measuring their activation by CD86 upregulation showed that only the 3F10 induced moDC activation (Fig. 7c). Furthermore, among the F4/80^+^ macrophage population, the percentages of both CD11c and MHCII positive macrophages increased with 3F10 treatment (Fig. 7d-f). This indicated maturation towards a DC-like phenotype, resembling the Ly6i^+^ MHCII^+^ CD11c^+^ population increased in the scRNA data set (Fig. 6a,b and S7f). Finally, macrophages from 3F10-treated mice also expressed higher levels of CD86 (Fig. 7g).

**Figure 7:**
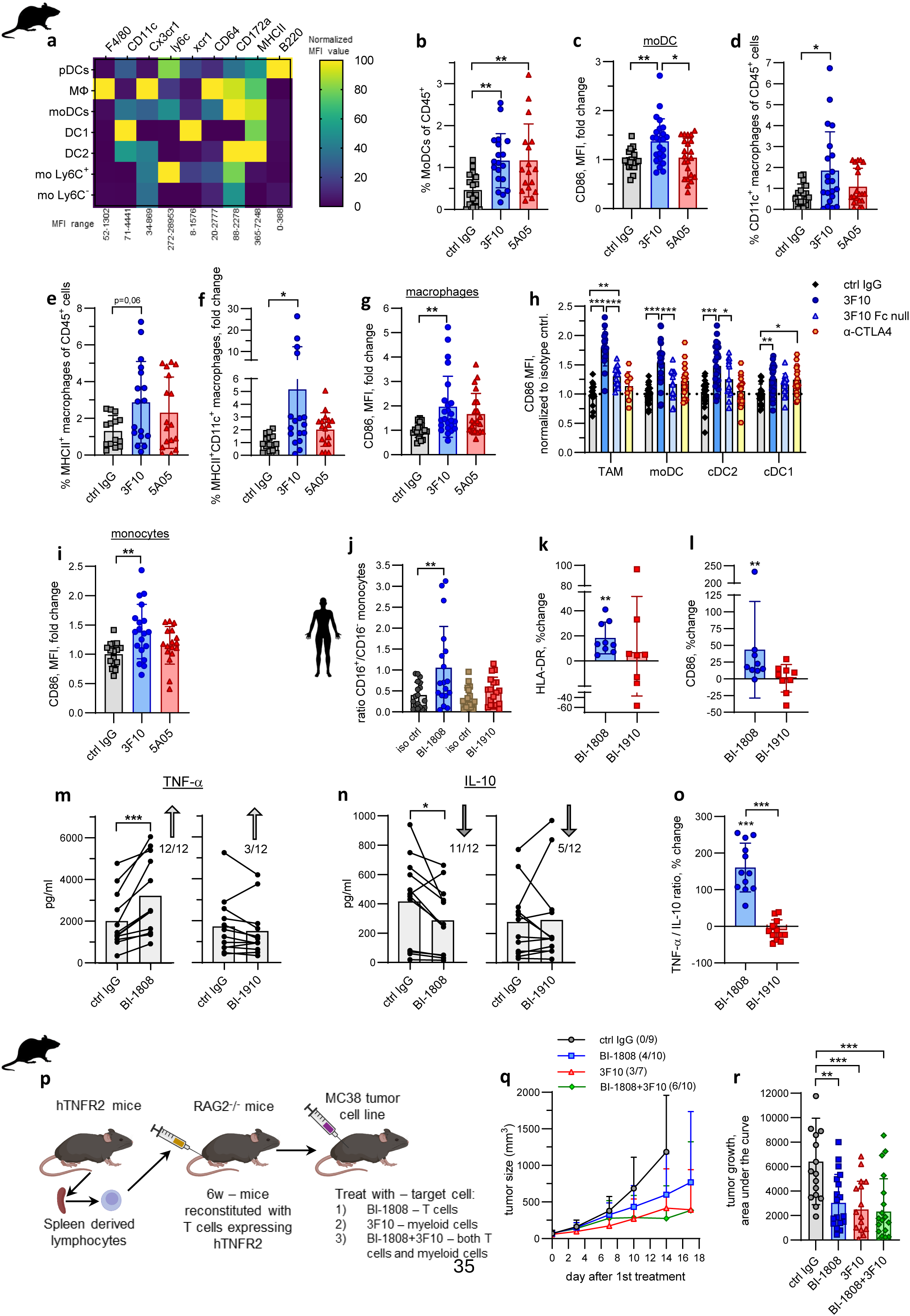
Myeloid reprogramming contributes to a-TNFR2 anti-tumor activity a-g and **i,** Shows data from cells isolated from tumors, day 8 after indicated treatment, analyzed by FCM. Data were collected from 3-5 independent experiments and are shown as the mean ± SD. If the baseline varied between experiments, the data were normalized to the control IgG and presented as the mean ± SD fold change. **a,** Heatmap of marker expression in the 7 different myeloid and DC cell types identified. The data was normalized where 100 corresponds to the highest, and 0 the lowest, MFI observed for that marker. The MFI range for each marker is shown below the heatmap. The data shown is from one representative experiment. **b,** Quantification of moDCs. **c,** CD86 expression in moDCs. **d-e,** Percent macrophages expressing CD11c or MHC II. **f,** Quantification of CD11c and MHCII double-positive macrophages. **g,** CD86 expression in macrophages. **h,** CD86 expression in macrophages and DCs isolated from CT26 tumors 6 h post first treatment. Data are from 3 independent experiments and show mean ± SD of values normalized to the mean of isotype control-treated mice in each experiment. Each dot represents one individual mouse. **i,** CD86 expression in tumor infiltrating monocytes. **j-n,** Human monocytes were isolated and incubated with antibodies as indicated for 24 h. Activation markers were analyzed using FCM, and cytokines using MSD or ELISA. Each dot represents one donor. Data were collected from 3 independent experiments. **j,** ratio of CD16^+^/CD16^-^ monocytes. **k-l,** HLA-DR and CD86 expression in treated CD16^+^ monocytes relative to isotype control. **m,** Change in TNF-a and **n,** IL-10 in supernatants from BI-1808 or BI-1910 treated, relative to control IgG treated, monocytic cultures. Arrows indicate the direction of change relative to isotype control, and numbers indicate in how many donors this change occurred. **o,** Ratio of TNF-a / IL10 production in monocytic cultured treated with BI-1808 or BI-1910. **p,** Schematic illustration of an experiment to create tumor-bearing mTNFR2:hTNFR2 chimeric mice where 3F10 treatment targets myeloid cells and BI-1808 treatment targets T cells. **q,** Tumor growth in chimeric mice treated as indicated, data from one representative experiment. Experiments were terminated when the first mouse/group reached experimental end-point due to tumor size. Number in parenthesis show numbers of tumor free / total number of mice at day 35. **r,** Accumulating tumor growth days 1-14 in chimeric mice as calculated using AUC for individual mice from 2 independent experiments. For statistics in b-g , i and r, one-way ANOVA was used, in j Repeated Measures one-way ANOVA was used, in h two-way ANOVA was used using Tukey’s adjustment for multiple comparisons, in k-n Student’s paired T-test was used, in o Student’s T-test was used. *=p<0,05, **=p<0,01 and ***=p<0,001.

To investigate whether the 3F10-mediated activation resulted from direct myeloid targeting or indirect effects mediated via CD8^+^ T cell activation, the experiment was repeated, this time terminating the mice 6 h after antibody treatment to reduce the likelihood of pronounced T cell activation. A strong upregulation of CD86 was detected on intratumoral myeloid cell populations also at this early time point (Fig. 7h). This effect largely depended on FcγR interactions and was diminished in mice treated with the 3F10 Fc-null variant. Further consistent with a direct myeloid targeting effect, no or very minor effects of α-CTLA-4 (which does not directly target myeloid cells) were observed. The majority of myeloid cells in a tumor originate from infiltrating monocytes^50,51^ and as shown in figure 1m-n, monocytes express high levels of TNFR2, facilitating direct signalling. Interestingly, 3F10 upregulated CD86 also on tumor-infiltrating monocytes (Fig. 7i). This data suggests that activation and reprogramming towards moDCs start already at the monocyte level.

For translation to the human system, we evaluated human monocyte polarization in vitro. In primary monocyte overnight cultures, ligand-blocking BI-1808 increased CD16 (FcγRIII) expression, causing a shift from CD16^-^ to CD16^+^ cells (Fig. 7j). Further studies showed that BI-1808 also increased HLA-DR and CD86 expression on the CD16^+^ monocyte population (Fig. 7k-l). Finally, provoking the monocyte cultures with a very low amount of LPS (20 pg/ml) to induce cytokine production showed that BI-1808 caused an increase in proinflammatory TNF-α which was parallelled by a decrease in anti-inflammatory IL-10 (Fig. 7m-o). None of these effects were seen with the purely agonistic antibody BI-1910. Together, these data indicate that BI-1808 and its surrogate antibody, 3F10, mediate similar myeloid remodeling, transitioning from tumor-supporting to pro-inflammatory phenotypes, which may be initiated in monocytes.

To address whether the myeloid reprogramming contributed to α-TNFR2’s anti-tumor effect, we created mouse/human TNFR2 chimeric mice by reconstituting RAG2^-^/^-^ mice with lymphocytes from human TNFR2 transgenic mice (for reconstitution, see Extended Data fig.8a-c). This resulted in mice expressing human TNFR2 on T cells and B cells and murine TNFR2 on myeloid and other cell types. Selective targeting of myeloid or T cell-expressed TNFR2, or both, was subsequently achieved using treatment with 3F10 or BI-1808 specific for mouse or human TNFR2, respectively (Fig. 7p). Tumor growth data strongly indicated that both myeloid and T cells contributed to α-TNFR2’s anti-tumor activity (Fig. 7q-r and Extended Data fig. S8d).

In summary, ligand-blocking α-TNFR2 induced profound reprogramming of myeloid cells in an FcγR-dependent manner, resulting in an increased number of activated MoDC contributing to tumor regression.

## Discussion

Existing literature supports a pleiotropic role for TNFR2 in immune regulation and highlights its potential as a target for cancer immunotherapy. Despite this, its pro- versus anti-inflammatory effects in different contexts and a lack of understanding of the desired antibody characteristics have hampered its therapeutic exploitation.

To address this and define the optimal antibody format, we generated two parallel sets of TNFR2 antibodies, targeting either human or mouse TNFR2, with both sets containing ligand blockers and non-ligand blocking agonists. Furthermore, we investigated isotype and FcγR dependence and demonstrated how the opposing properties of the two sets of antibodies can both elicit robust CD8^+^ T cell mediated anti-tumor immunity across multiple tumor models.

Ligand-blocking antibodies 3F10 and BI-1808, reduced intratumoral T regs, dependent upon activating FcγRs. This was presumably through antibody-dependent cellular cytotoxicity or phagocytosis, as described for other T reg-expressed TNFRSF members ^41,52–54^ as well as CD25 and CTLA-4 ^55–57^. Interestingly, only the CCR8^+^ T reg population was depleted, and the reason for this is currently under investigation. This population has previously been described as the most suppressive T regs associated with tissue-homing and intratumoral clonal expansion ^58–62^. Consequently, several CCR8 targeting antibodies are under development for cancer treatment ^63–65^. It is well established that T regs suppress CD8^+^ T cell-mediated immunity ^66,67^, and multiple reports indicate that activating FcγR-mediated T reg depletion contributes to clinically relevant effects of the α-CTLA-4 and α-TIGIT mAbs ^48,56,68–70^. Based on the known impacts of T reg depletion, it is likely that this contributes to tumor regression ^63,64^.

Unexpectedly, our MoA studies showed that engagement of the inhibitory FcγRII was also required for maximal therapy of the ligand-blocking antibodies (Fig. 2a-d). This contrasts with observations from T reg-depleting antibodies to other TNFR receptors, e.g., GITR ^71^, OX40 ^42,52,54^, and with CTLA-4 ^56^, where single-agent activity relies solely on activating FcγRs, strongly suggesting 3F10’s mechanism-of action extends beyond pure T reg depletion. Conversely, it also contrasts with agonist antibodies to CD40, whose in vivo efficacy depends on FcγRII but not activating FcγRs ^72, 39,73^. Antibodies targeting 41BB or OX40 can mediate depletion or agonism dependent on antibody isotype ^41,42,54^. However, these activities were shown to be competitive, requiring sequential administration of T reg-depleting and CD8 T cell-promoting variants for optimal therapeutic effects. Data presented here suggests that ligand-blocking α-TNFR2’s dual and simultaneous engagement of both activating and inhibitory FcγRs is linked to its profound remodeling of the myeloid system. While the TME in control-treated mice was dominated by tumor-promoting M2-like macrophages, the α-TNFR2 antibody evoked a potent switch to M1-like macrophages and moDCs. Using a chimeric mouse model where myeloid and T cells were separately targeted (Fig. 7p-r), we showed that myeloid-restricted TNFR2-targeting also induced anti-tumoral effects. Recent publications show analogous, albeit less pronounced, FcγR-mediated transformation of intratumoral macrophages to M1-like phenotypes using α-CTLA-4 ^48^ or α-TIGIT antibodies ^69^. Translational observations indicate the clinical relevance of this, highlighting FcγR-mediated myeloid reprogramming as an important and previously overlooked requirement for successful immunotherapy ^55,69,74^. Interestingly, for α-CTLA-4 and α-TIGIT, the efficacy and macrophage reprogramming effects required a high A:I FcγR binding format ^43,48,69^, while the ligand-blocking α-TNFR2 mAb could utilize either activating or inhibitory FcγRs and achieved superior therapeutic efficacy when engaging both. Given the heterogenous FcγR expression in tumors^75^, the ability to engage any FcγR type to elicit anti-tumor immunity may be critical. A major difference between TNFR2 and CTLA-4 or TIGIT is their expression pattern. CTLA-4 and TIGIT are primarily T cell targets ^76,77^, but TNFR2 is additionally highly expressed on myeloid cells. Importantly, myeloid cells express the full repertoire of FcγRs, which can mediate receptor crosslinking and clustering of TNFR2 and thereby signaling ^39,73,78^, whilst concurrently triggering ITAM activation (through the activating FcγR) in the myeloid cell ^79–81^. This would allow α-TNFR2 to directly target the myeloid cells, which could account for the profound re-programming effects observed here. Mechanistically, the observed T reg depletion and myeloid reprogramming, together with the dual FcγR requirements and differentiated target expression, suggest that BI-1808 achieves something that has not previously been observed in a single TNFRSF targeting antibody, combining mechanisms and effects seen with e.g. α-GITR (T reg depletion) and α-CD40 (myeloid agonism). This understanding is important for the development of therapeutically optimal α-TNFR2 drugs, especially considering the T reg focus currently dominating the TNFR2 field. For example, recent work has suggested improved T reg targeting by designing an FcγRIIIA-enhanced TNFR2xCCR8 bispecifics ^27^. However, based on the mechanistic characterization herein of ligand-blocking α-TNFR2, such an antibody will likely have far less effect on myeloid cells and more limited therapeutic potential.

The clearest difference in anti-tumor efficacy between the two different TNFR2 antibodies was seen in the immune-excluded B16 model, where the ligand-blocking antibodies outperformed the agonists. We hypothesize that the powerful myeloid reprogramming evoked by the ligand-blockers results in a local inflammation transforming the TME to facilitate immune cell recruitment and subsequent anti-tumor response, without which the agonists cannot induce potent effects. Thirdly, a strong tumor-specific CD8^+^ T cell activation is achieved both intratumorally and in tumor-draining lymph nodes with the ligand-blocking TNFR2 mAb. Whether the CD8^+^ T cell activation is achieved through antibody-dependent co-stimulation, because of T reg depletion, myeloid activation, or a combination thereof, remains to be determined. Regardless, it is instrumental for this antibody’s anti-tumor activity, with efficacy ablated in the absence of CD8^+^ T cells.

The agonistic antibody had little impact on the myeloid compartment but activated all TNFR2-expressing lymphocytes, confirming several earlier publications describing TNFR2 as a costimulatory molecule important for amplifying and sustaining T cell responses ^28–31,38^. Consistent with previous findings with TNFRSF agonists ^39,73,78^, the agonist α-TNFR2 antibody showed intrinsic co-stimulatory activity enhanced by inhibitory FcγRII engagement. Importantly, lymphoid cells are high in TNFR2 but low in TNFR1 ^82^. Both receptors exist as cell-surface and soluble variants and undergo negative feedback loops of self-regulation where TNF-α signaling results in an immediate release of soluble TNFR2, acting as a decoy receptor ^83–85^. Based on the observed differences in lymphocyte versus myeloid agonism between the antibodies, one can hypothesize a model where the receptors compete for a limited amount of ligand. In this scenario, a non-ligand-blocking agonistic antibody would preferentially agonize cells with high TNFR2 and simultaneously low ligand-competing TNFR1 expression. Investigating the TNFR2/TNFR1 expression ratio of immune cells in the CT26 tumor model (Fig S9) supports this hypothesis as the highest ratio is observed on T cells and NK cells, which are broadly activated by the agonist. Translation of this knowledge to therapeutic development could result in combinations with a pure T reg-depleting agent to avoid unwanted effects of T reg agonism. Alternatively, one of the major challenges with TIL or CART cell therapy for solid cancer is the lack of sustained T cell expansion, and combinations with a TNFR2 agonist could potentially overcome this ^86–88^. Finally, although the instrumental role of CD8^+^ T cells in anti-tumor immunity is widely demonstrated, increasing amount of data highlights the importance of CD4^+^ T cells in supporting tertiary lymphoid structures ^89,90^ and sustaining CD8^+^ T cell responses ^91,92^- here the effects of a TNFR2 agonist like BI-1910 remains to be explored .

Despite clear differences in their early mechanisms of action, both α-TNFR2 mAbs evoked a profound activation of tumor-specific CD8^+^ T cells and tumor rejection, supporting TNFR2 as a powerful new immunomodulatory target for cancer treatment. Moreover, both mAbs combine effectively with checkpoint blockade, overcoming resistance to α-PD-1, positioning them well as a new paradigm for treatment and efficacy in combination studies.

Given the two antibodies divergent initial mechanisms of action, it will be interesting to explore in what scenarios each mAb may be more effective. Ultimately, the importance of each will only be realized in vivo in human trials. To support clinical translation, our results were verified using two human lead candidates: the first-in-class ligand-blocker BI-1808 and the intrinsic agonist BI-1910, both assessed in vitro and in a transgenic mouse expressing human TNFR2. These lead candidates are currently in clinical development for solid cancer, and we are excited to confirm their principal mechanisms of action in humans and see their impact on tumor control.

## Supporting information

Martensson et al Supplemental Figures and Tables

## Material and Methods

### Generation of antibodies targeting human or mouse TNFR2

Human and mouse extracellular domain of TNFR2 (target protein) were purchased from Sino Biological Inc, both including a C-terminal polyhistidine tag (His-tag). CD40 from Sino Biological Inc was used as non-target protein. The TNFR2 proteins were biotinylated with ChromaLink^TM^ Biotin (SoluLink) according to standard procedures. Antibody fragments against TNFR2 were isolated from the proprietary human phage display library n-CoDeR^®^. Enrichment of specific TNFR2 antibodies was achieved by three consecutive pannings using either the biotinylated TNFR2 protein or TNFR2 coated to polystyrene balls. The amount of target used decreased with each panning. A pre-selection with 1) 200 pmol of the biotinylated non-target (human or mouse CD40) loaded on Dynabeads, 2) naked Dynabeads and 3) 500 pmol of the non-target protein coated in immunotubes (Nunc) was done prior to each panning. In all pannings, bound phages were eluted using trypsin and analyzed by colony forming unit (CFU) titration to estimate the number of phages in eluted phage pools. Prior to use as input in the next panning eluted phage pools were amplified in *E. coli*.

The different phage pools were analyzed by ELISA and flow cytometry to evaluate the enrichment of target-specific binders. For ELISA, phage stocks were titrated in a two-fold serial dilution starting at 2×10^11^ CFU/ml in plates coated with either target (2pmol/well), non-target (1pmol/well) or streptavidin (0,1µg/well) and left to bind for 3 h at room temperature. Bound phages were detected using anti-M13-HRP (GE Healthcare) followed by OPD (Sigma Aldrich) addition and absorbance reading at 490/650nm.

For flow cytometric analysis, phage stocks from each panning were added at 1:4, 1:16 and 1:64 dilutions to hTNFR2 or mTNFR2 or mock transfected 293-FT cells. Anti-M13 (GE Healthcare) was added after 2 h incubation followed by anti-mouse-APC (Jackson Immunoresearch) detection.

### Samples were analysed in a HTFC Screening system (Intellicyt)

Single-chain fragments from panning 3 were converted from phage-bound to soluble format and expressed as individual clones in *E. coli*. TNFR2 transfected cells were seeded into FMAT plates. *E. coli* expressed scFv were added followed by mouse anti-HIS antibody (R&D Systems) and anti-mouse-APC (Jackson Immunoresearch). The assay was performed without washing steps. Stained cells were detected using FMAT (8200 cellular detection system, Applied Biosystems). Positive scFv clones were picked, re-expressed and re-tested for binding by ELISA and on transfected cells using flow cytometry, as described above. For ELISA TNFR2 protein was coated in ELISA plates at 2 pmol/well. Supernatant of *E. coli* expressed scFv was added and left to bind for 1 h at room temperature. Bound antibodies were detected using anti-FLAG-AP (Sigma Aldrich) followed by substrate addition (CDP-star, Life Technologies) and luminescence reading (Tecan Ultra).

ScFv that specifically bound TNFR2 were sequenced over CDR H1, H2 and H3 using the Sanger method. Unique hTNFR2 targeting clones were converted to human IgG1 while mTNFR2 targeting clones were converted to mouse IgG2a as previously described^1^.

### Ligand-blocking ELISAs

Purified human TNFR2 or mouse TNFR2 (Sino Biological) were coated to assay plates at 50 nM/well. α-TNFR2 mAbs were added at 67 nM and left to bind for 1 h at room temperature followed by addition of human TNF-α-bio (R&D Systems) at 5 nM or mouse TNF-α (Gibco) at 2 nM. Ligand was left to bind for 15 minutes before detection by Streptavidin- HRP (Jackson ImmunoResearch) or rabbit-anti-TNF-α (Sino Biological) plus anti-rabbit-HRP (Jackson ImmunoResearch) followed by substrate addition (Pierce, Thermo Fischer) and luminescence reading (Tecan Ultra). For BI-1808 and 3F10, the assay was repeated with titrated concentrations of antibodies.

### Target cross-reactivity ELISAs

Human TNFR1-His, TNFR2-His or LTBR-His (Sino Biological) were coated overnight at 10-20 nM/well in assay plates. After washing, antibodies were added in titrated doses starting at 67 nM and left to bind for 1 hour at room temperature. Bound antibodies were detected using anti-human F(ab)-HRP (Jackson Immunoresearch) followed by substrate addition (Pierce, Thermo Fischer) and luminescence reading (Tecan Spark). The same assay was performed for mouse surrogates but using purified mouse TNFR1-hFc-His (R&D Systems), TNFR2-His (Sino Biological) or LTBR-hFc (Abcam) coated at 1,5-10 nM. Antibody detection was done using anti-mouse F(ab)-HRP (Jackson Immunoresearch).

### Human IL-10 and TNF-α concentration in monocytic culture supernatants

The concentration of IL-10 and TNF-α present in culture supernatants of isolated monocytes incubated in presence of BI-1808 was quantified using sandwich immunoassay kits (R&D Systems, DY210 and DY217B) according to manufacturer’s instructions. Briefly, supernatants from overnight cultures were diluted and added to microplates previously coated with capture antibodies against the target cytokine.

After washing the wells, detection antibodies were added to the plate followed by another washing step and incubation with HRP-streptavidin solution for signal amplification. Substrate solution was added immediately before reading optical density at 450 nm using a Tecan Spark Microplate reader. Standard curves for the respective cytokine targets were used to interpolate cytokine concentrations in GraphPad Prism v10.5.0 for Windows (GraphPad Software), and dilution factors were included in the final concentration values before further analysis.

### Generation of cell lines

All cell lines were subcultured at 37 °C with 5% CO_2_. CT26 was cultured in complete RPMI (Glutamax supplemented with 10% FBS, 1 mM sodium pyruvate and 10 mM HEPES). NK-925 GFP-CD16 cells were cultured in MEM alpha with 10% FBS, 10% Horse serum, 0,1 mM b-Mercapto-ethanol (Sigma), 0,2 mM Myo-Inositol (Sigma), 2,5 nM folic acid (Sigma), 1x Non-essential amino acids, 1 mM sodium pyruvate, 1x Penicillin/Streptomycin and 10-100 IU/ml rh-IL2 (R&D systems). All reagents from Invitrogen unless stated otherwise.

To generate TNFR1 and/or TNFR2 deficient cell lines, 5×10^5^ cells of either CT26 (ATCC) or NK-925 (Conquest) were electroporated with ribonucleoparticles (RNPs) consisting of Alt-R CAS9 enzyme, Alt-R CRISPR-Cas9 crRNA and Alt-R CRISPR-Cas9 tracrRNA according to manufacturer protocols (Integrated DNA Technologies). NK-925 cells were electroporated with 1820V, 20 ms and one pulse; CT26 cells were electroporated with 1300V, 20 ms and 2 pulses. After 2-3 days of recovery, cells were used to generate monoclonal cell lines using the limited dilution method. After 2 weeks of incubation, monoclonal cells were amplified and then analyzed by FCM for lack of expression of the target protein. Knock-out of the respective protein was confirmed by Sanger sequencing.

To generate Jurkat reporter cells the human TNFR2 gene (NM_001066) was inserted into pcDNA3.1(+) and this plasmid was used for transfection of NF-κB/Jurkat/GFP™ Transcriptional Reporter Cell Line (Systems Biosciences, Cat nr TR850A-1). 5 x 10^6^ cells were incubated with 30 μg plasmid DNA on ice for 10 min whereafter the cells were pulsed using BioRad electroporator (settings = 0.3 kV, 960 μFD) and incubated on ice for further 10 mins. The cells were then cultured in complete media (RPMI with 10% FCS, 2 mM glutamine, 1 mM pyruvate) in 96-well plates. Two days post transfection selection geneticin (ThermoFisher Scientific, cat 10131027) at final concentration of 1 mg/ml was added and cells were incubated for 2-3 weeks until individual colonies developed. After establishing the cell lines, 1×10^5^ cells were plated in a flat-bottom 96-well plate with or without addition of 1×10^5^ CHO-K1 mCD32b cells, generated as described before^2^. BI-1808 or BI-1910 was added to the cell culture and incubated for 6 hours at 37 °C and 5% CO_2_. The % GFP^+^ Jurkat cells were analysed by FCM, samples were acquired on FACS Canto II (BD).

### Western blot, NF-κb signaling

TNFR1^-^/^-^ NK-925 cells were incubated with indicated antibodies for 15 min. Thereafter cells were lysed in RIPA buffer supplemented with phosphatase and protease inhibitor cocktails (Thermo Scientific). Protein extracts were collected after centrifugation at 16500 x g for 15 min and 4 °C. Quantification was performed using Bradford Protein Assay kit (Thermo scientific) and thereafter the lysates were complemented with 4x sample buffer (LI-COR) and reducing agent (Thermo Scientific) together with heat treatment at 95 °C for 5 min. Protein lysates were separated with 50 ug per lane on a 7.5% Criterion™ TGX™ Precast Gel (Bio-Rad) and blotted onto a Immobilon-FL PVDF Membrane (Millipore). Membranes were dried for 1 h at room temperature and thereafter the protein loading was analyzed using Revert 700 Total Protein stain (LI-COR). After blocking, the membranes were incubated with primary antibodies against IκBα (1:750), NF-κB p65/RelA (1:1.000 and β-Actin 1:10. 000, all antibodies from Cell Signaling Technology) at 4 °C overnight, followed by 1 h at room temperature with secondary antibodies (IRDye® 800CW Goat anti-Mouse IgG and IRDye® 680RD Goat anti-Rabbit IgG, both LI-COR). Finally, an Odyssey Fc from LI-COR was used for visualization and Empiria Studio for analysis of the results.

### NK cell assay

The NK cell assay used has been previously described by Almishri et al. ^3^. In brief, human NK cells were isolated from human PBMCs by MACS (NK cell isolation kit, Miltenyi). 1×10^5^ NK cells were added per well in NK medium (RPMI + Glutamax (Invitrogen), 10% HI FBS (Invitrogen), 2 mM L-glutamine, 50 µM 2-mercaptoethanol, 100 μg/ml Penicillin/Streptomycin (all Gibco)) and cultured with 20 ng/ml rhIL-2 and 20 ng/ml rhIL-12 together with 10 μg/ml α-TNFR2 mAb, 10 μg/ml ctrl IgG or 100 ng/ml rhTNF-α (all cytokines were from R&D Systems). Supernatants were collected after 24 h and the amount of IFN-γ produced was assessed using V-plex Human IFN-γ Kit (Meso Scale Diagnostics (MSD)) and read by QuickPlex SQ 120 Reader (MSD). Data were analyzed using Discovery Workbench v.4.0 software (MSD).

### In vitro activation of CD4^+^ T cells

Human PBMCs were isolated from buffy coats (Halmstad hospital, Sweden) using Ficoll Paque PLUS gradients (Cytiva). CD4^+^ T cells were isolated by MACS (CD4^+^ T cell isolation kit (human), Miltenyi). Mouse CD4^+^ T cells were isolated from spleens by MACS (CD4^+^ T cell isolation kit (mouse), Miltenyi). CD4^+^ T cells were cultured in R10 medium (RPMI + Glutamax (Invitrogen), 10% HI FBS (Invitrogen), 1 mM sodium pyruvate, 10 mM hepes, 100 µg/ml Penicillin/Streptomycin (all Gibco)) and activated with 50 ng/ml rhIL-2 or 135 IU/ml rmIL-2 (both R&D Systems). For binding studies, “Dynabeads Human T-Activator CD3/CD28 beads” or “Dynabeads Mouse T-Activator CD3/CD28 beads” (both ThermoFisher) were added at 1:1 ratio for 48-72 h. Cultured cells were labelled with viability dye and antibodies detecting CD4 and CD25 (see FCM panel table 1). The α-TNFR2 mAbs were added at concentrations ranging from 0.002-267 nM. For T cell activation assays, 1×10^5^ purified human T cells were stimulated with plate-bound anti-CD3 (0.6 µg/ml; clone UCHT1, R&D Systems) plus 10 µg/ml of soluble, cross-linked or not cross-linked α-TNFR2 mAb and 1 µg/ml anti-CD28 (clone CD28.2, BioLegend) for 72 h at 37 °C and 5% CO_2_. To cross-link, antibodies were incubated with F(ab’)_2_ goat anti-human IgG, Fcg fragment specific (Jackson ImmunoResearch), in a molar ratio IgG:F(ab’)_2_ of 1.5:1 for 1h at room temperature before addition to cultures. Cells were washed and stained with viability dye and antibodies against CD4, CD8 and CD25 (see FCM panel table 2). Cells were acquired on FACS Verse (BD). All FCM data were analyzed with FlowJo software (Tree Star).

### Tumor associated macrophages (TAM) : T cell co-cultures

Human TAM- like cells (hereafter TAMs) were generated by culturing MACS-isolated, human CD14^+^ monocytes (CD14^+^ microbeads, Miltenyi) in 50% sterile filtered ascitic fluid isolated from cancer patients, and 50% R10 medium for 72 h at 37 °C and 5% CO_2_. TAMs were then washed and pre-incubated with 10 µg/ml α-TNFR2 mAb for 30 min. Without washing the cells, titrating numbers of TAMs were co-cultured with MACS-isolated (pan T cell isolation kit, Miltenyi), CFSE labelled (2 µM, Invitrogen) CD3^+^ T cells in the presence of Dynabeads (above) in a 1:3 (bead:cell) ratio. After 72 hours, the cells were stained with viability dye and CD25 antibody (see FCM panel table 3).

Mouse TAMs were isolated from digested CT26 tumors (described) using MACS (CD11b^+^ microbeads, Miltenyi). Naïve T cells were isolated from Balb/C mice spleens by MACS (pan T cell isolation kit, Miltenyi) and labelled with 2 µM CFSE (Invitrogen). TAMs were plated and pre-incubated with 10 μg/ml α-TNFR2 mAb for 30 min. Without washing the cells, titrating numbers of TAMs were co-cultured with T cells in the presence of “Dynabeads Mouse T-Activator CD3/CD28 beads” (ThermoFisher) in a 1:3 (bead:cell) ratio. After 72 h, the cells were stained with viability dye and CD25 antibody (see FCM panel table 3).

Cultured cells were acquired using FACS Verse (BD).

### Mice

Six to eight weeks-old (17-20 g) female C57BL/6J and BALB/c were either purchased from Janvier (Saint-Berthevin, France) and maintained at BioInvent local facilities or purchased from Charles River (Kent, UK) and bred and maintained in UK Home Office approved local facilities. C57BL/6J mice expressing human TNFR2 in the place of murine TNFR2 were licensed from Biocytogen (Beijing, China) and breed at BioInvent facilities. Fcγ-chain KO BALB/c, CD32KO and γ-KO/CD32KO BALB/c mice are described in^4^ , and bred in UK Home Office approved local facilities. The genotype and phenotype status of the mice was confirmed by either PCR/flow cytometry.

Mouse/hTNFR2 chimeric mice were created by reconstituting B6Rag2 mice with cells from C57BL/6J mice expressing hTNFR2. Spleens from hTNFR2 transgenic mice were homogenized into a single cell suspension through a 100 μm cell strainer and depleted from CD11b and CD11c positive cells by MACS cell separation (Miltenyi Biotec). 5 x 10^6^ cells per mouse were injected iv into B6Rag2 mice.

Reconstitution was confirmed in blood 4 weeks post injection. Blood was stained with viability dye and antibodies against CD45, CD4, CD8 and TCRβ (see FCM panel table 4). T cells and classical monocytes (CD11b^+^Ly6C^high^) expression of human and mTNFR2 in chimeric mice were established using FCM. Blood was drawn from vena saphena (BD Vacutainer Heparin tubes) from chimeric, C57BL/6J and hTNFR2 mice and TNFR2 expression was obtained using biotinylated BI-1808 (human) and 3F10 (mouse). Biotinylation of antibodies was done using Chromalink (Trlinkbio) according to manufacturer’s instructions. For further staining details, see FCM panel table 4. Cells were analyzed on FACSLyric (BD).

For in vivo experiments performed at BioInvent, all mice were housed and maintained in accordance with Swedish Laboratory Animal Care guidelines and were performed in an animal care-accredited facility. All experimental animal studies were conducted under the approval of the Swedish Ethical Committee (approval numbers 5.8.18-17196/2018 and 5.8.18-19686/2022) and carried out in accordance with the Federation of Laboratory Animal Science Associations (FELASA) guidelines. For in vivo experiments performed at University of Southampton, all mice were housed in IVC units and maintained in accordance with the Animal (Scientific Procedures) Act 1986 and experiments were performed under approved project licenses (P4D9C89EA and PP5396109), and ethical review by local University of Southampton committee (approval numbers 69840, 65400).

### In vivo mouse tumor models

The CT26, EMT-6, B16-F10 (B16) and EG7 murine carcinoma cell lines were obtained from ATCC. The MC38 cell line was kindly gifted by Dr Sjef Verbeek. Cells were cultured according to providers instructions and logarithmic growth phase of cells was ensured before harvesting cells for grafting. A total of 0.5 x 10^6^ or 1 × 10^6^ CT26, B16 or MC38 cells were inoculated subcutaneously into the flank of each mouse. For EMT-6 0.5 x 10^6^ tumor cells/mouse were injected into the mammary fat pad. All tumor cells were inoculated in 100 μl of PBS. After 8–13 days, mice bearing tumors of 40-200 mm^3^ (depending on model and setting) were randomized into treatment groups based on tumor size. For tumor growth inhibition studies mice were treated with the following antibodies or antibody combinations. Anti-mouse PD- 1 (29F.1A12, rat IgG2a, 10 mg/kg, BioXcell), anti-mouse CTLA-4 (9H10, hamster IgG 10 mg/kg, BioXcell), anti-TNFR2 (3F10 mIgG2a, mIgG1 and mIgG1-N297A; 5A05 mIgG2a, mIgG1 and mIgG1-N297A; BI-1808 hIgG1, BI-1910 hIgG2 and/or control antibodies mIgG2a, mIgG1 or hIgG1 isotype). Antibodies were administered intraperitoneally in a volume of 200 μl at 10 mg/kg (Balb/c) or 40 mg/kg (C57BL/6J) due to different PK-profiles in the different mouse strains. For CD8 depletion experiments, mice were pretreated with 1 mg of YTS169-mIgG2a (produced in-house) one day prior to tumor inoculation and four days after tumor inoculation. For NK-cell depletion 10 µL was anti-Asialo-GM1-pIgG (BioLegend, cat: 146002) diluted in 200 µL PBS on days -2, 4 and 11. Depletion was confirmed by screening blood or spleen by FCM, see FCM panel table 5.

Animals were continuously monitored, and mice were euthanized when any of the following end points were met: study termination, tumor burden ≥2,000 mm^3^, tumor ulceration, or moribund appearance. Tumor burden was measured using calipers, and tumor volumes were calculated using the modified ellipsoid formula 0,5 × (length × width^2^).

In immune profiling analysis and scRNA-seq experiments, antibodies were administered in 1 or 3 treatments 3-4 days apart. To identify mice responding to treatment, blood was drawn from vena saphena (BD Vacutainer Heparin tubes) day 1 and 7 post treatment for biomarker analysis. Then, 24-48 h after treatment, mice were euthanized and tumors, tumor draining lymph nodes and spleens were collected.

Tumors were diced and incubated with either 1 mg/mL of collagenase Type V (Merck) in a total volume of 1 mL PBS; then incubated at 37 °C at 250 rpm for 20 minutes or in a solution of 50% Liberase (Roche) and 50% DNAse (Roche) for 3x5 min with slight vortexing in between at 37 °C, 5% CO_2_. Digestion reaction was quenched with 5 mL of RPMI (containing 10% FCS) and homogenized into a single cell suspension through a 100 μm cell strainer. Lymph nodes and spleen did not undergo enzymatic digestion but were homogenized into a single cell suspension through a 100 μm cell strainer. Red blood cells in the spleen were lysed using Lysis Buffer (Gibco).

### Cell preparation for single cell RNA sequencing

Tumors harvested as indicated were enzymatically digested as described. Viable lymphocytes were isolated using gradients (Lympholyte-M Separation Media, Cedarlane). Tumor lymphocytes were stained with viability dye and CD45 antibody (see FCM panel table 6) and viable CD45^+^ cells were sorted using FACS Aria Fusion (BD). Cells were counted (NucleoCounter NC-202, ChemoMetec) and samples were fixed using 4% paraformaldehyde (PFA) for 20 hours at 4 °C, followed by long-term storage at –80 °C according to ‘10x Genomics’ recommendations. Samples were further processed by Center for Translational Genomics at the Medical faculty at Lund University, Sweden. In short, prior to library preparation, samples were thawed, washed, and hybridized with barcoded probe sets according to the Chromium Next-GEM Flex protocol (10x Genomics). Approximately 10^4^ cells per sample were targeted for capture, and the pooled samples were loaded onto a Chromium X instrument using a Chip Q. Libraries were sequenced on an Illumina NovaSeq 6000, targeting a sequencing depth of 2×10^4^ reads per cell.

Animals responding to treatment day 8 were identified on day 7 by measuring the tumor volume and an increase of CD8^+^ effector memory T cells and tumor-specific CD8^+^ T cells using MHC class I Dextramers (H-2Ld – SPSYVYHQF-APC (AH-1), Immudex). The whole blood was stained with viability dye and antibodies against CD62L, CD45, CX3CR1, CD4, CD44, CD8 and TCRβ (see FCM panel table 7). Red blood cells were lysed using BD FACS Lysing Solution (BD) before acquired on LSRFortessa X-20 (BD).

### Single cell RNA analyses

The scRNA-seq reads were aligned to the mouse transcriptome and UMI counts are measured to generate an expression matrix using the Cell Ranger Pipeline. The output files from cellranger were converted to a Seurat object using the R Seurat package (v4.4.0) for downstream analysis ^5^. The R package Doubletfinder (v2.0.3) was applied with default parameters to identify and remove cell doublets in each sample individually ^6^. The cell cycle phase was controlled with the Seurat *CellCycleScoring* function using cell cycle related genes. The low-quality cells were removed based on the number of detected genes, the number of detected UMIs, house-keeping gene expression and the percentage of mitochondrial gene expression. The samples were normalized using the *SCTransform* function and applied separately for each dataset after finding the top 3000 feature genes using *SelectIntegrationFeatures* function. As a next step, we applied functions *PrepSCTIntegration*, *FindIntegrationAnchors* to find integration anchors and *IntegrateData* (normalization.method = "SCT") to complete data integration.

For clustering of single cells, we run principal component analysis on the gene space using *RunPCA* and clustering was performed based on the shared nearest neighbor between cells (*FindNeighbors*) and graph-based clustering (*FindClusters*). We used uniform manifold approximation and project (UMAP) with the function *RunUMAP* with the same number of principal components used for embedding for the visualization of clusters. For the final clustering, dimensional reduction using UMAP used the top 30 calculated PC dimensions and a resolution of 0.6, eventually identified 26 distinct immune cell subpopulations.

To identify marker genes, we used *FindAllMarkers* function in Seurat (one-sided Wilcoxon rank sum test) with p-value adjusted for multiple testing using the Bonferroni correction. We used curated cell specific marker genes identified and based on prior experience to annotate the immune cell subpopulations. For cell-type specific sub-clustering, we integrated the data across samples and performed *SCTransform* approach as mentioned above with parameters ranging from 20-30 PCs and UMAP resolution ranging between 0.2–0.7 with final parameters selected to generate consistent visualizations.

*AverageExpression* function in Seurat was used to calculate the average expression of gene markers and z-scores per genes were calculated for visualization with the pheatmap package. The R package LSD (https://cran.r-project.org/web/packages/LSD/index.html) was used to visualize density plots of cells using the UMAP coordinates of cells from each condition. To visualize mRNA expression of selected marker genes, we used R Schex package (v1.14.0) and converted the UMAP embedding into hexbin (nbin=40) quantifications of the proportion of cells that express indicated marker gene.

The cell trajectory analysis was performed using the R package Slingshot (v2.8.0) with default settings using the umap cell embeddings calculated in seurat as the input^7^

### Characterization of immune cell contents in tumor mouse models analyzed by FCM

WT mice were challenged with either CT26, EMT6 (BALB/c) or MC38 (C57BL/6J) and tumors were monitored for growth. When CT26 tumors reached approx. 600-700 mm^3^ or MC38/EMT6 reached approx. 1300-1400 mm^3^ in area, mice were sacrificed, and samples were processed as described above. Immune cell populations were identified using the FCM panels in table 14; the proportion of each cell population was calculated as a percentage of total CD45.2^+^ cells.

### Tumor-infiltrating and tumor-draining LN immune cells analyzed by FCM

Single cells from tumors and tumor-draining LNs were obtained as described at 6 h, day 2 and day 8 post first treatment. Cells were stained with 4 different FCM panels (see FCM panel table 7 - 10). For intracellular markers, cells were fixed and permeabilized using the Foxp3/Transcription Factor Staining Buffer Set (Invitrogen). Cells were analyzed on LSRFortessa X-20 (BD). For gating strategy, see Supplementary Fig. 1-3.

### Tumor antigen-specific CD8^+^ T cells

Tumors and spleens were harvested from CT26 mice 10 days after the first treatment and prepared according to above. 1 x 10^6^ isolated tumor cells were restimulated with 2 μg/ml of tumor specific peptides (AH-1, SPSYVYHQF, BioNordika). Tumor cells were pulsed for 4 h in the presence of brefeldin A (Sigma) at 37 °C and 5% CO_2_. Isolated splenocytes were restimulated for 48 h at 37 °C and 5% CO_2_, the last 4 h in presence of brefeldin A (Sigma). Cytokine-producing CD8^+^ T cells were then identified by FCM staining for CD45, TCRβ, CD8, TNF-α, IFN-γ and CD25 (see FCM panel table 13). For intracellular markers, cells were fixed and treated as described above. In addition, tumor-specific CD8^+^ T cells in CT26 and MC38 tumors were identified using MHC class I Dextramers (CT26: H-2Ld – SPSYVYHQF-APC (AH-1), MC38: H- 2 Kb/KSPWFTTL-APC (p15E) and H-2 Db/ASMTNMELM-APC (Adpgk), Immudex; see FCM panel table 7). Cells were analyzed using LSRFortessa X-20 (BD). For gating strategy, see Supplementary Fig. 1.

### Tumor-infiltrating myeloid cell FCM analysis

For day 8 data, raw FCS files were subjected to FlowAI plugin with default settings to remove bad events before analysis. For heat-mpa generation, clean files originating from the two groups (isotype control and 3F10) were then gated for CD45^+^ lineage negative (TCRb, CD19, NK1.1, Ly6G, SiglecF and CD49b) live single cells and concatenated in a single file with all events included, see FCM panel table 11. The proportion of each cell population was calculated as a percentage of total Live CD45^+^ lineage negative cells. Cells were analyzed using LSRFortessa X-20 (BD). For gating strategy day 8, see Supplementary Fig. 2a-b. For the 6 h time point gating strategy, see Supplementary Fig 3 and FCM panel 12.

### TNFR2 and TNFR1 expression assessment in naïve and tumor-bearing mice

CT26 tumor-bearing mice were culled, and tumors collected when approximately 250 mm^3^. Blood was collected from the tail vein, both from tumor bearing mice and age-matched naïve, into tubes containing 10% heparin and then kept on ice until staining. 10^5^ cells (or 10 μL whole blood) were stained with extracellular cell population markers for 30 minutes at 4 °C in FACS tubes. Red blood cells were lysed with 1 mL of Red cell lysing buffer (Bio-Rad cat # BUF04C) for 3-5 minutes at room temperature before staining. For intracellular staining, cells were fixed with fixation buffer overnight at 4 °C, and permeabilized with permeabilization buffer (both ThermoFisher Scientific). 1 μL/sample of FoxP3 was added and opsonized at 4 °C for 1 hour before washing and acquisition on FACS Canto II. Antibody panels used are described in table 8. Each sample was stained for the same cell populations, but either TNFR1 or TNFR2 were included. The MFI of the whole cell population (in the PE channel) was taken for both the TNFR1 and TNFR2 stained samples. These values were then used to determine the ratio of TNFR2/TNFR1 relative expression

### TNFR2 expression assessment in naïve human blood cells

Whole blood collected from healthy human subjects (in BD Vacutainer EDTA tubes). Cells were stained with antibodies panels listed in the FCM panel table 14. For intracellular markers, cells were fixed and permeabilized using the Foxp3/Transcription Factor Staining Buffer Set (Invitrogen). Cells were analyzed on LSRFortessa X-20 (BD).

### Human monocytes cultures

Leucocyte concentrates were obtained from healthy donors from Halmstad Blodcentral, Sweden. To isolate PBMCs, concentrates were diluted with PBS (Invitrogen) and added to previously prepared LeucoSep tubes (Greiner) with Ficoll-Paque PLUS (Cytiva). After centrifugation at 1200 x g for 20 min at room temperature, PBMCs were aspirated from mononuclear cell layer in a Ficoll gradient. CD14^+^ monocytes were isolated from 20×10^6^ PBMCs using anti-human CD14 microbeads (Miltenyi Biotec) according to manufacturer’s protocol. 2×10^5^ isolated monocytes were cultured per well with 10 µg/mL of TNFR2 mAb or isotype control in RPMI 1640+Glutamax (Invitrogen) and 10 % FCS for 20 h at 37 °C and 5 % CO_2_. For cytokine induction, 20 pg/ml LPS was added to cultures at the same time as antibodies. Supernatants were collected and concentrations of IL-10 and TNF-α were assessed. Cells were stained with viability dye and antibodies for surface markers CD14, CD16, HLA-DR, CD86 and CD11b (see FCM panel 15). For gating strategy see Supplementary Fig. 2c. Monocyte characterization by FCM was performed in a LSRFortessa X-20 (BD).

### TNFR2 blocking experiments

In vitro activated mouse splenic T cells (described in previous section) blocked with 10 μg/ml polyclonal TNFR2 (R&D), ctrl IgG, 3F10 or 5A05 for 30 min were stained with 2.5 μg/ml biotinylated 3F10/5A05 for 30 min. Cells were subsequently stained with viability dye, anti-CD25 and streptavidin. Incubations were done at 4°C. Cells were analyzed using a FACS Verse (BD) and the capacity of 3F10/5A05 binding to CD25^+^ cells after various blocking conditions analyzed.

### Statistical analysis

All data points in a set of experiments were included in statistical calculations. Outliers were only removed if there was a clear technical error in data sampling. Raw data were used when possible. For data sets with larger interexperimental variations in base line, (due to e.g. different settings in voltage of a flow cytometer), each data point was normalized to the mean value of the control IgG in the respective experiment, to enable pooling of several experiments. ScRNA data processing has been described above and no statistical analysis were performed in this data set. Data distribution was examined and mean±SD were used when appropriate.

For comparisons between two groups with normal distribution, Student’s T test was used. If the data points in each group originated from the same donor, paired Student’s T test was used. For comparisons regarding one parameter using three or more groups with normal distribution, one-way ANOVA test was used. If the data points in each group originated from the same donor, Repeated Measures ANOVA test was used. For comparisons regarding one parameter but over several e.g. cell types using three or more groups with normal distribution, two-way ANOVA test using Tukey’s adjustment for multiple comparisons was used. For survival curves, Log-Rank Mantel Cox test was used.

Number of data points, number of experiments, graphic interpretation (mean and SD etc.), units, statistical test used and p-value limits have been stated in figure legends. Individual data points were shown when possible.

No missing data tests were used.

---

**Extended Data fig. 1:**
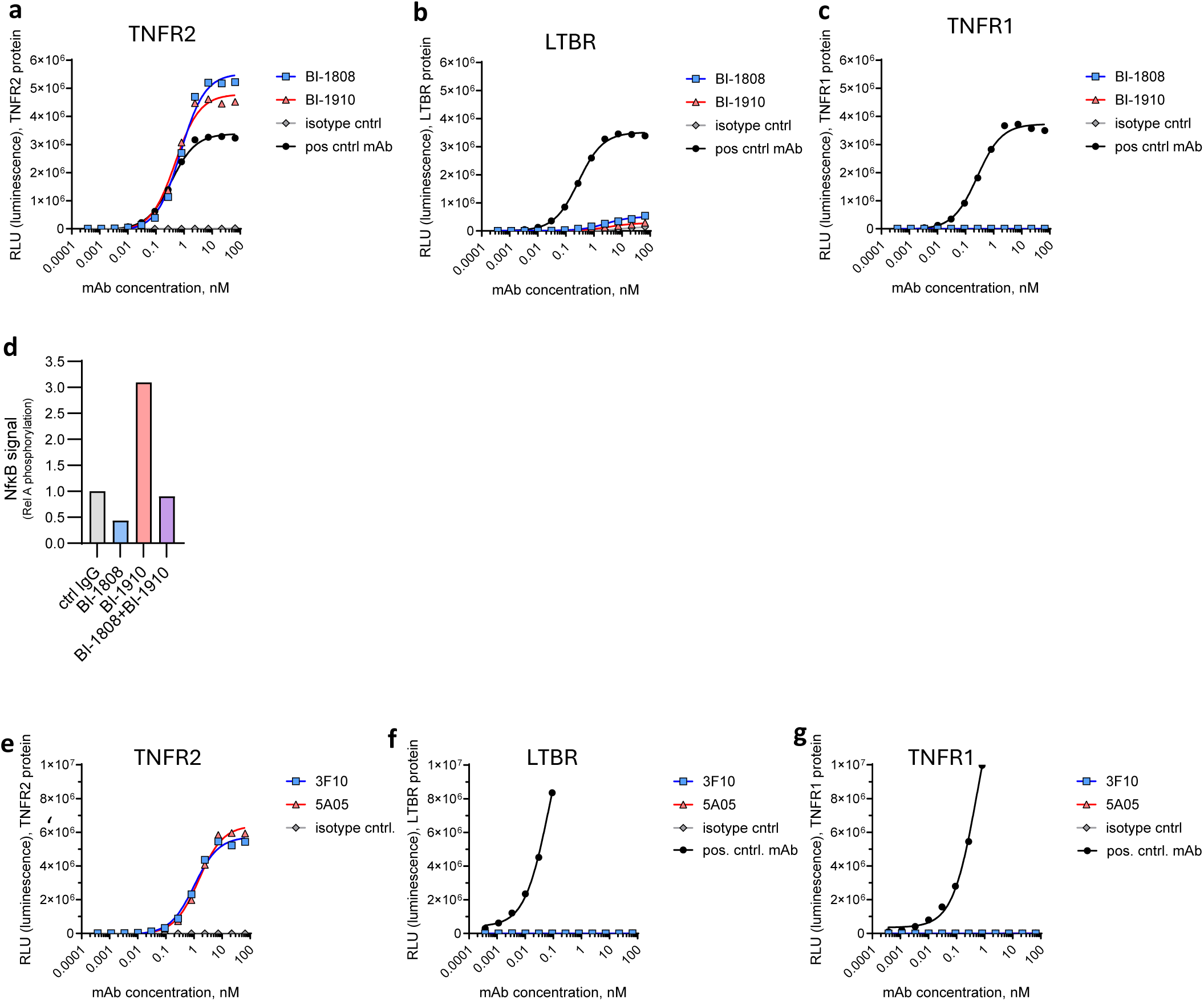
Generated antibodies are TNFR2 specific. BI-1808 and BI-1910 binding to **a,** TNFR2, **b,** LTBR and **c,** TNFR1 assessed by ELISA. **d,** NF-kB activation in TNFR1^-/-^ cells treated with 10 ng/ml TNF-a. Results shown from one representative experiment. **e-g,** 3F10 and 5A05 binding to **e,** TNFR2, **f,** LTBR and **g,** TNFR1 assessed by ELISA.

**Extended Data fig. 2:**
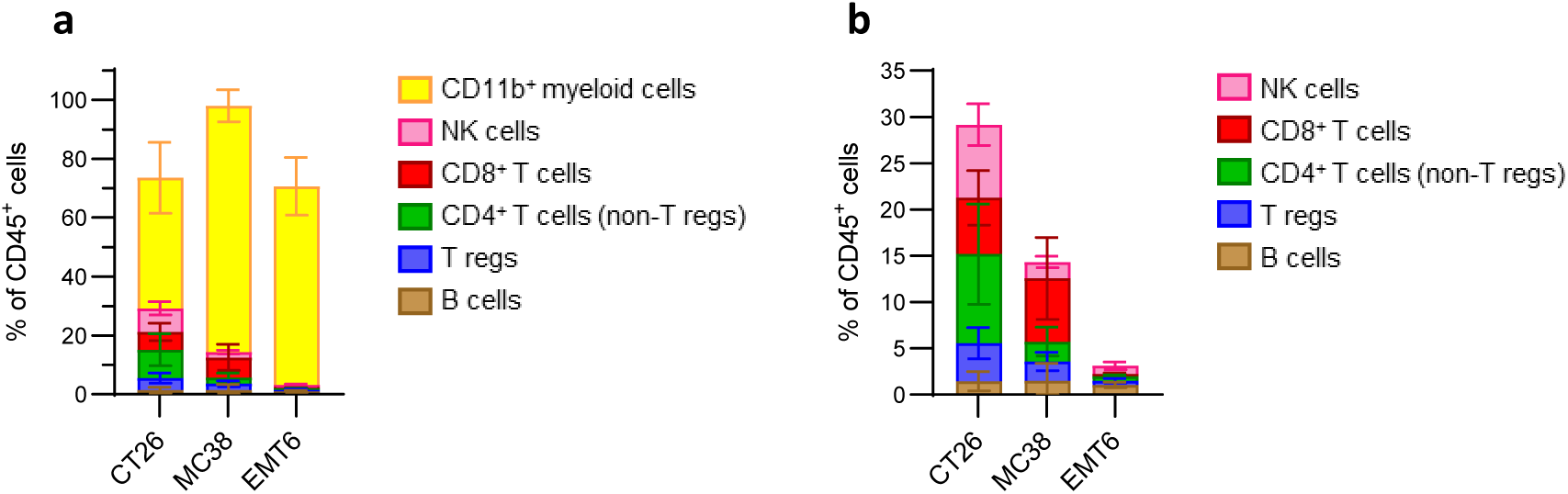
Immune cell infiltration in CT26, MC38 and EMT6 mouse models. a, Immune cell composition of CT26, MC38 or EMT6 tumors at sizes between 10×10 and 14×14 mm. **b,** Percentage of T, B and NK cells of total CD45^+^ cells from CT26, MC38 or EMT6 tumors at the same time points.

**Extended Data fig. 3:**
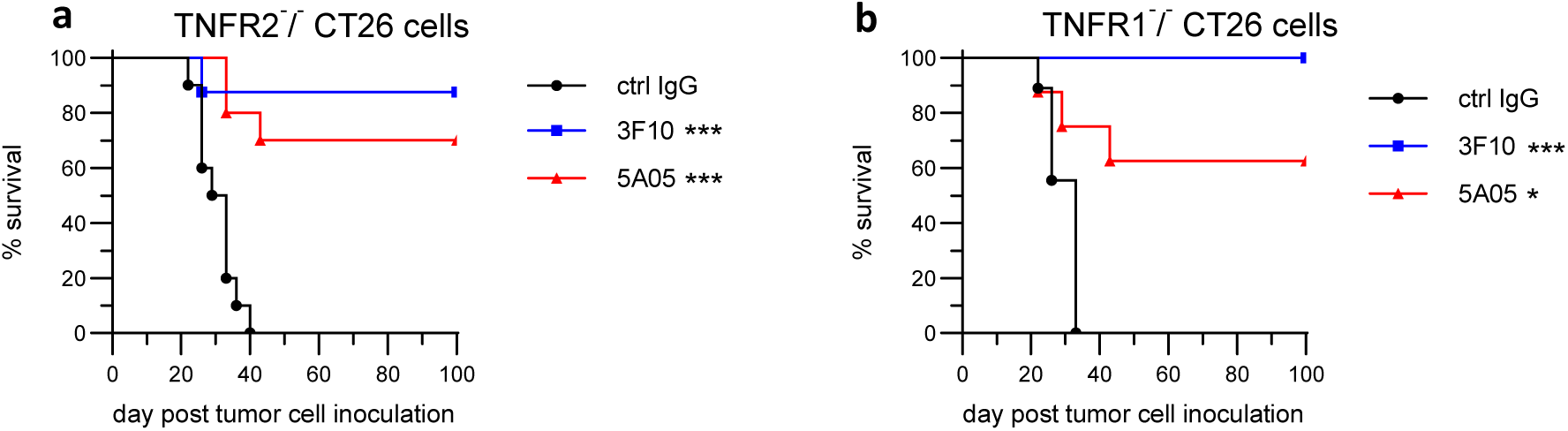
Anti-tumor effect is independent of tumor TNFR2 or TNFR1 expression. Survival curves in mice inoculated with **a,** TNFR2^-^/^-^ and **b,** TNFR1^-^/^-^ CT26 cells. ***=p<0,001 and *= p<0,05 using Log-rank (Mantel-Cox) test.

**Extended Data fig. 4:**
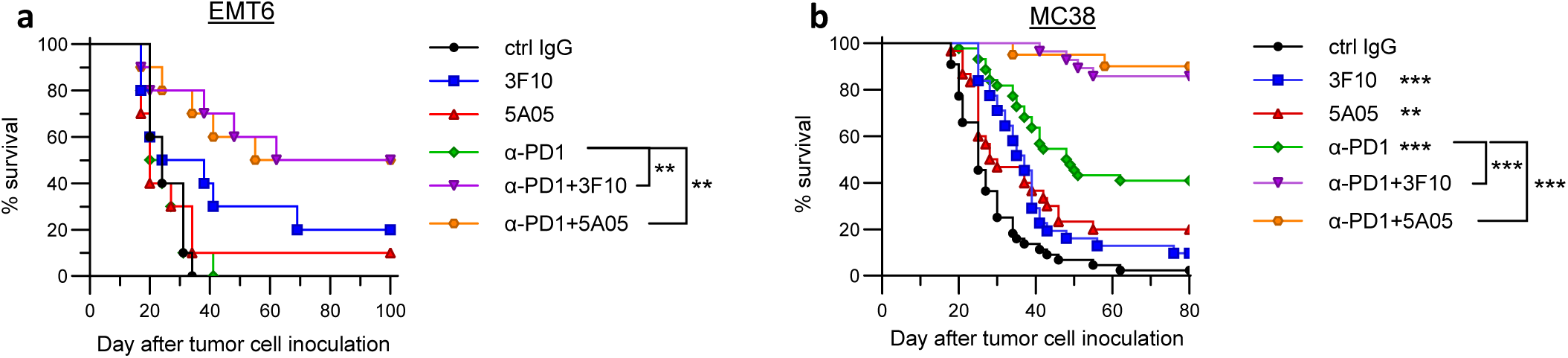
Both types of a-TNFR2 antibodies have broad anti-tumor effect and synergize with a-PD1. a-c, Survival of tumor-bearing mice treated with 3F10 or 5A05, as single agent or in combination with a-PD-1, showing **a,** data from one representative experiment (N=10 mice/group) when tumors were 100-200 mm^3^ at the start of treatment. **b,** Data from five different experiments (N=20-45 mice/group) when tumors had an average size of 140 mm^3^ at the start of treatment and the a-TNFR2 antibodies were used at a suboptimal dose..

**Extended Data fig. 5:**
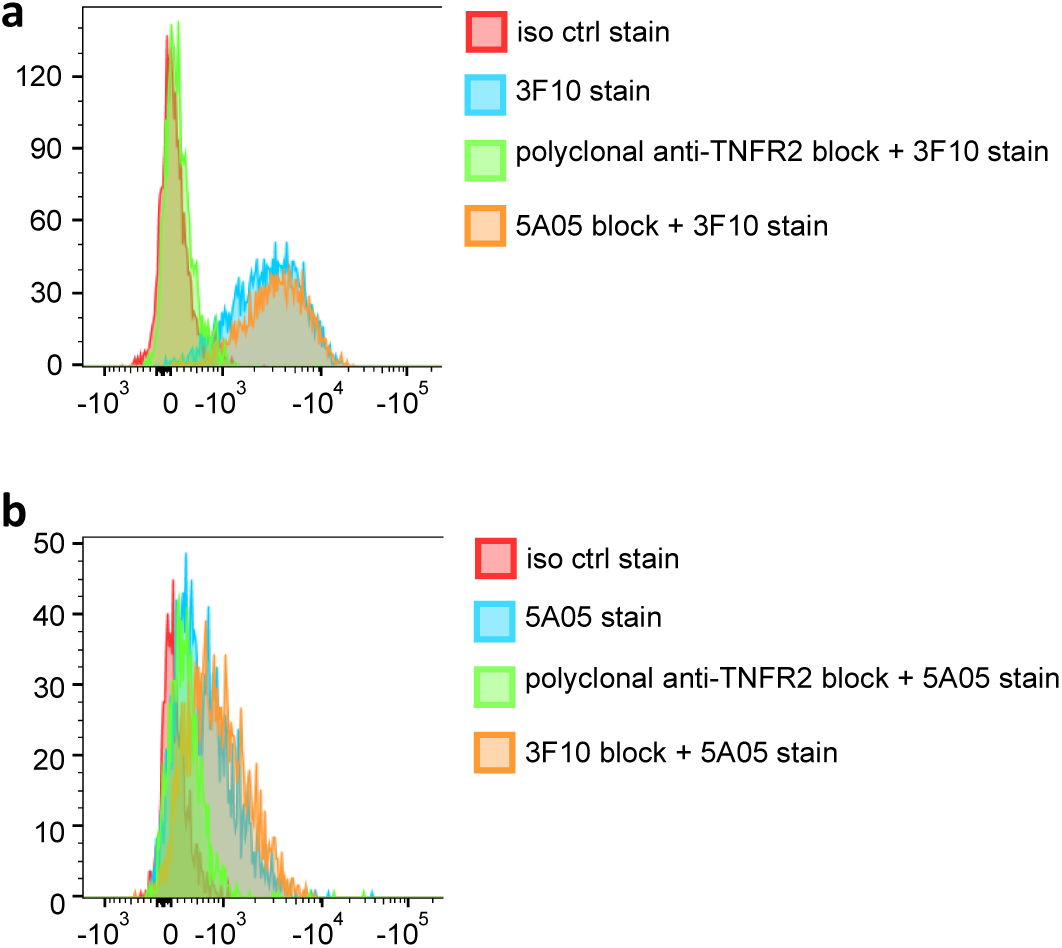
Ligand-blocking 3F10 and agonist 5A05 bind independently of each other a, FCM study on in vitro activated T cells preincubated with antibodies as indicated and stained with biotinylated 3F10 antibody. **b,** FCM study on in vitro activated T cells preincubated with antibodies as indicated and stained with biotinylated 5A05 antibody.

**Extended Data fig. 6:**
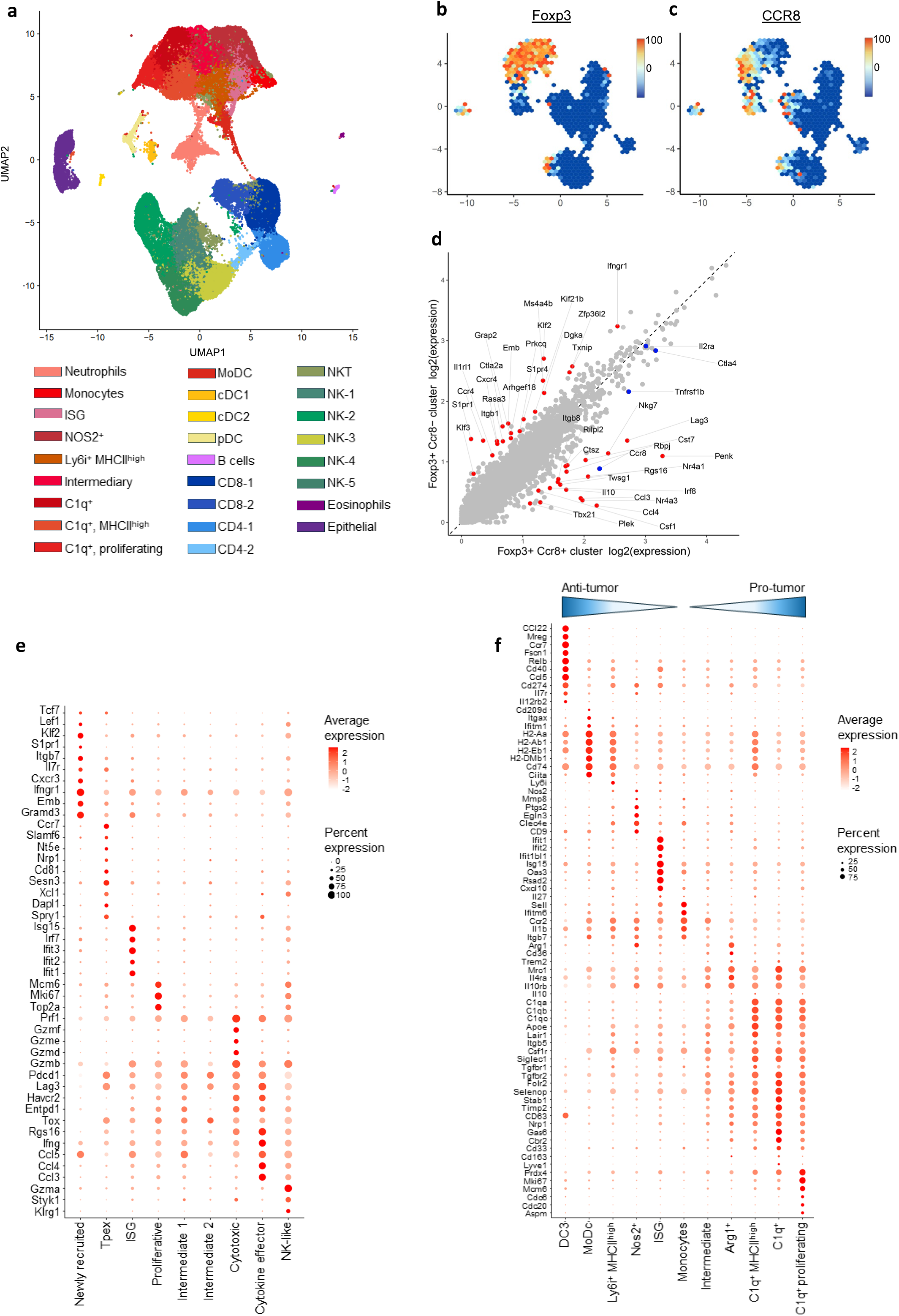
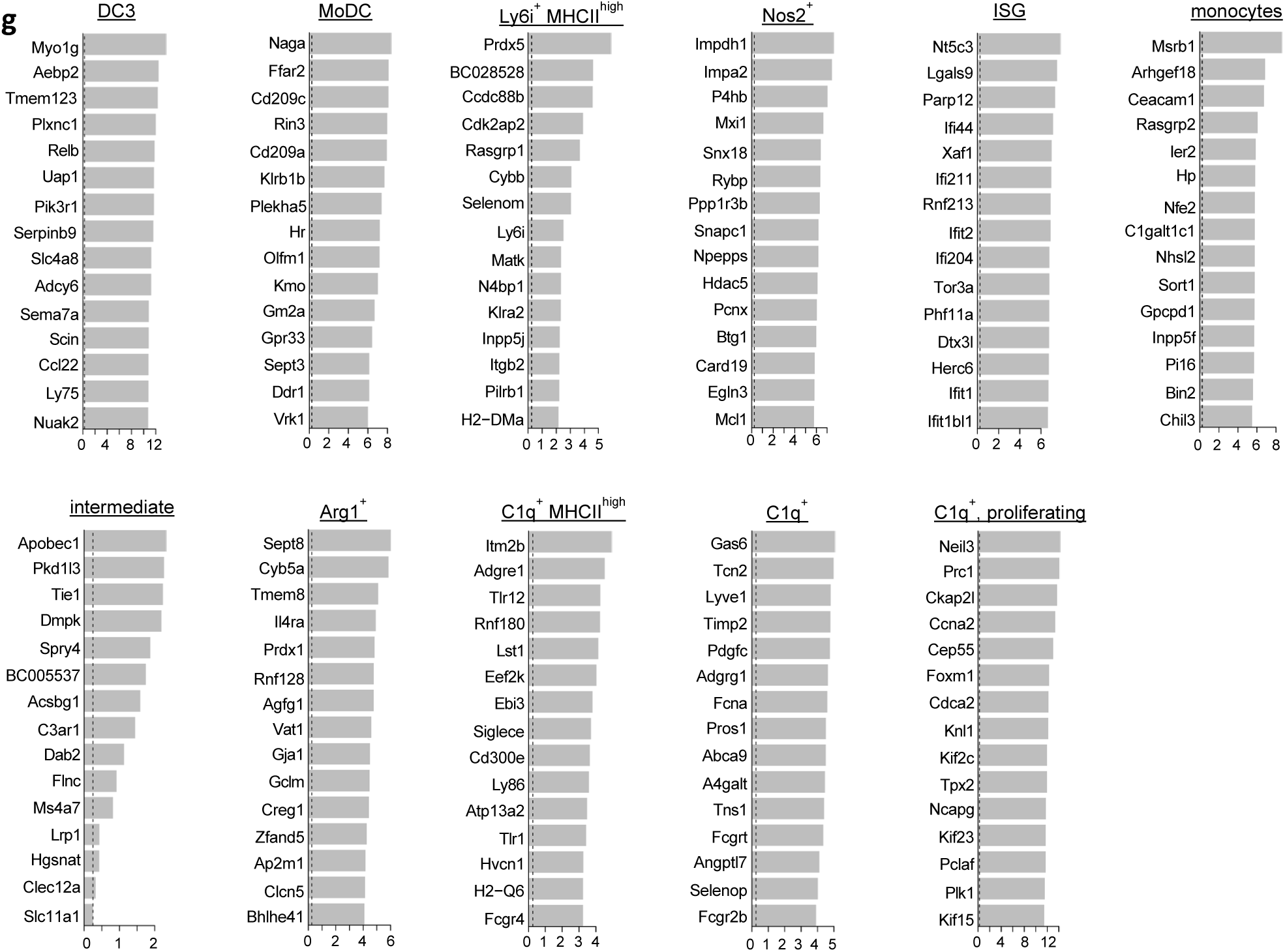
**Characterization of transcriptionally distinct cell populations following antibody treatments**. **a,**UMAP clustering of 63482 cells colored by immune cell subset from 10 tumor samples originating from 5 treatments (ctrl IgG, 3F10, 5A05, a-CTLA-4 and a-PD-1). Each group contains 10 mice and two time points (days 2 and 8 post first treatment, 5 mice/group and time point). **b,** Expression of *Foxp3* in the CD4^+^ T cell cluster. **c,** Expression of *CCR8* in the CD4^+^ T cell cluster. Red indicates high expression, and blue indicates low expression. **d,** Scatterplot shows differentially expressed genes comparing the CCR8^+^ and CCR8^-^ Foxp3^+^ CD4^+^ T cell subclusters. The top differently expressed genes (log2Fold change >1 and FDR <0.05) are highlighted in red. *Ctla4, CCR8, Il2ra* and *Tnfrsf1b* genes are highlighted in blue. **e,** Bubble plot showing marker gene expression across CD8^+^ T cells. **f,** Bubble plot showing marker gene expression across myeloid cells to define the different cell populations. The dot size indicates the fraction of expressing cells colored according to average normalized expression levels. **g,** Top 15 differentially expressed genes for the myeloid cell sub-clustering populations. The x-axes indicate log2 fold-change compared to all other cell populations

**Extended Data fig. 7:**
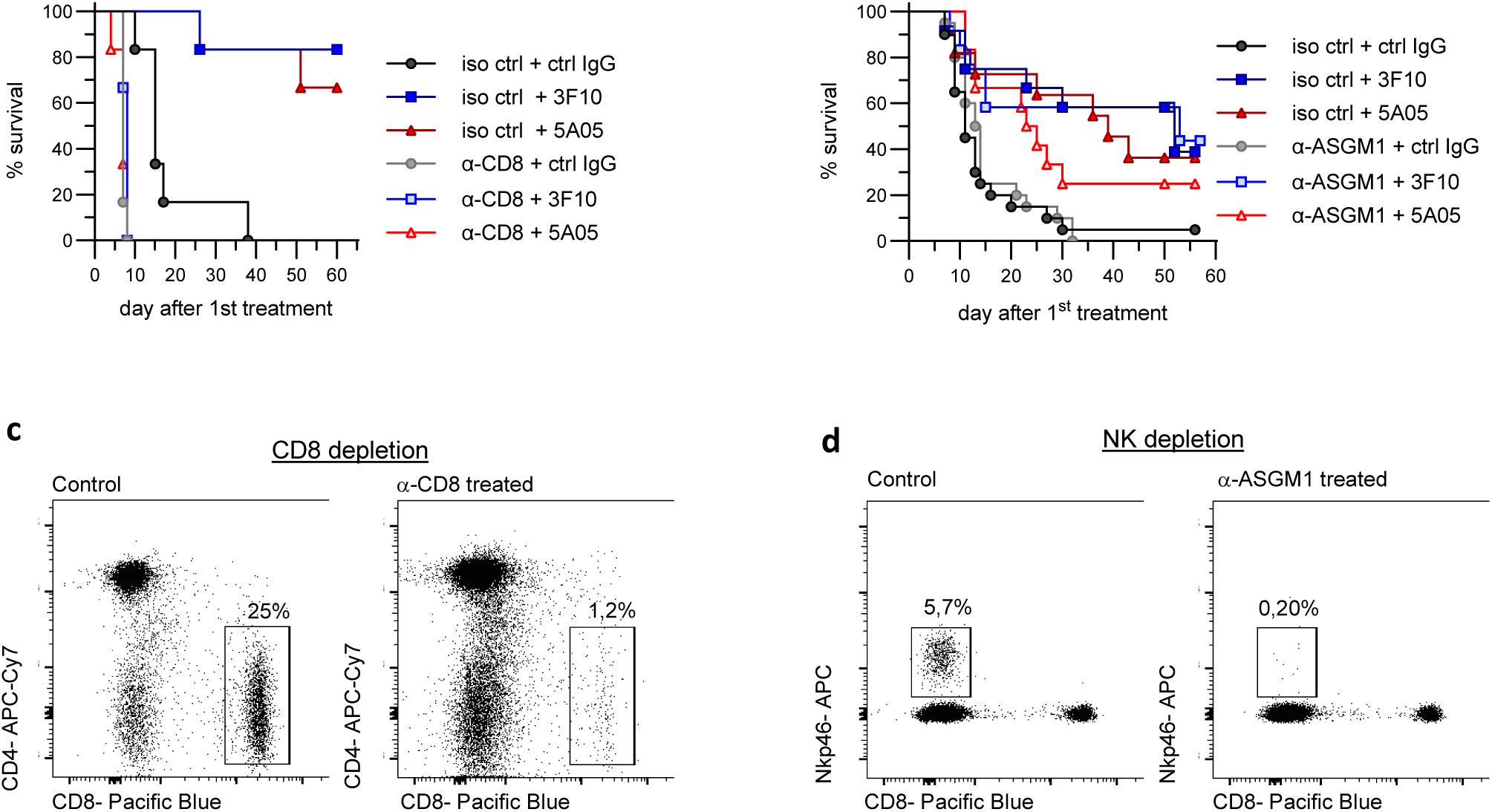
CD8 and NK cell depletion a, Survival curves of CT26 tumor bearing mice depleted of CD8^+^ cells prior to antibody treatment. **b,** Survival curves of CT26 tumor bearing mice depleted of NK cells prior to antibody treatment. **c,** Depletion of CD8^+^ cells confirmed by FCM. **d,** Depletion NK cells confirmed by FCM.

**Extended Data fig. 8:**
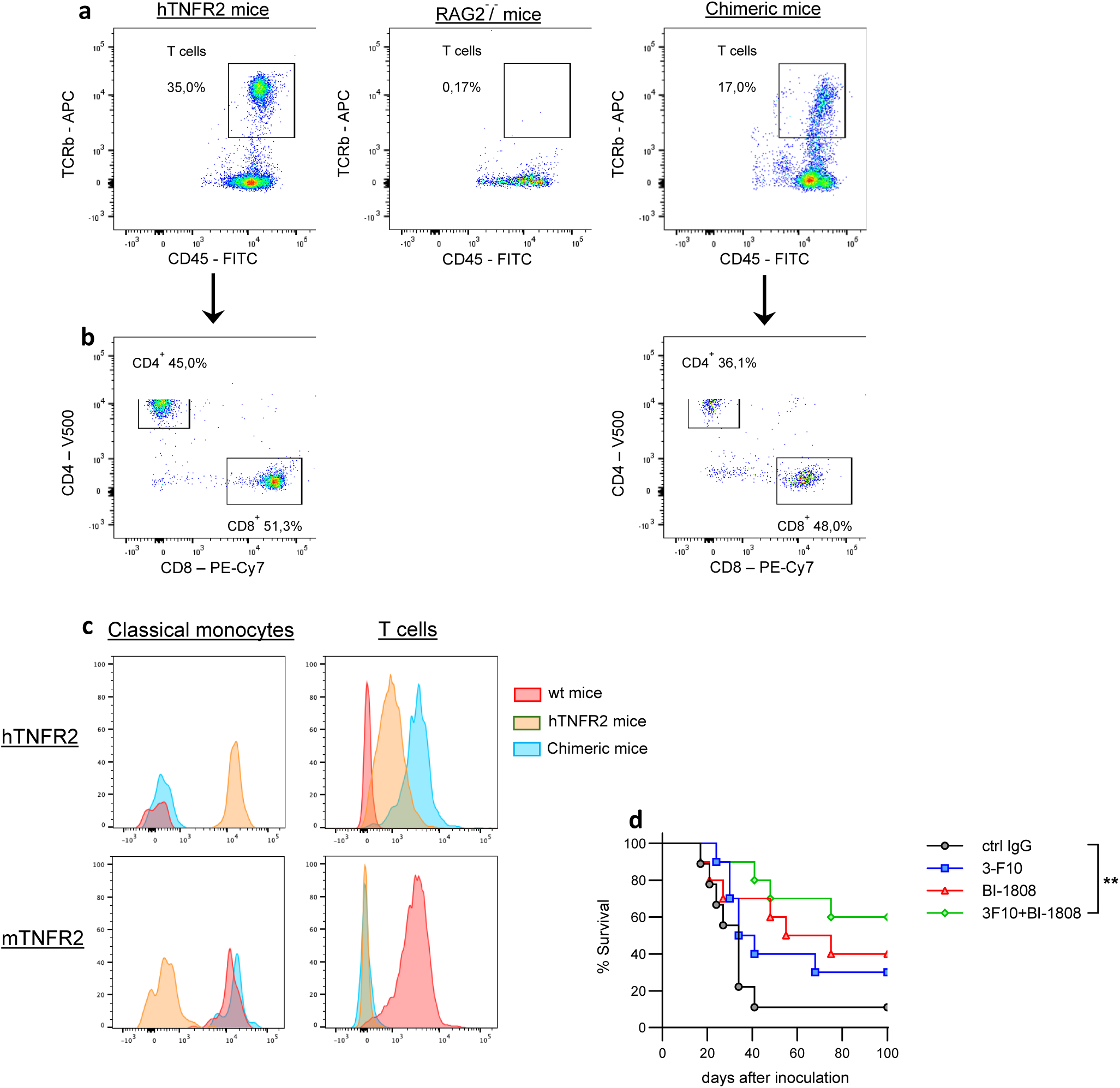
**RAG^-^/^-^ mouse reconstitution with hTNFR2^+/+^ lymphocytes**. **a,** Representative mice from each group showing normal T cell levels in hTNFR2 transgenic mice; RAG2^-^/^-^ mice lacking T cells; and chimeric mice reconstituted with lymphocytes from hTNFR2 mice. T cells are quantified by double staining for CD45 and T cell receptor b-chain. **b,** CD4^+^ and CD8^+^ T cell distribution in hTNFR2 mice and chimeric reconstituted mice. **c,** Expression of human TNFR2 and mouse TNFR2 on classical monocytes and T cells in chimeric, C57BL/6J (wt), and hTNFR2 mice. **d,** Survival curves in MC38 tumor-bearing chimeric mice treated as indicated. **=p<0,01 and Log-rank (Mantel-Cox) test.

**Extended Data fig. 9:**
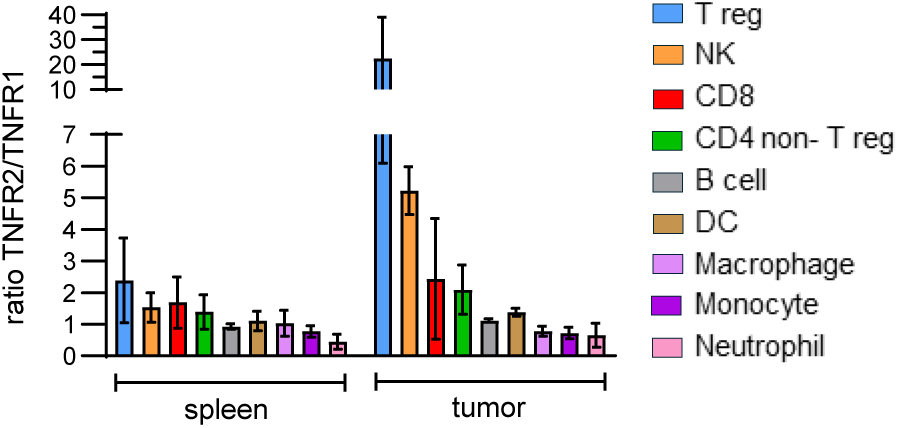
TNFR2 / TNFR1 ratios, CT26 tumor-bearing mice. TNFR1 and TNFR2 expressions on CD45^+^ immune cells were analyzed using FCM in CT26 tumor-bearing mice in spleen and tumor and an MFI based TNFR2/TNFR1 ratio was calculated.

