## Supplementary material for "Leveraging TNFR2 for antitumour immunity: T reg depletion and myeloid reprogramming versus T cell costimulation": Martensson et al Supplemental Figures and Tables

### 1 Supplementary figures and tables

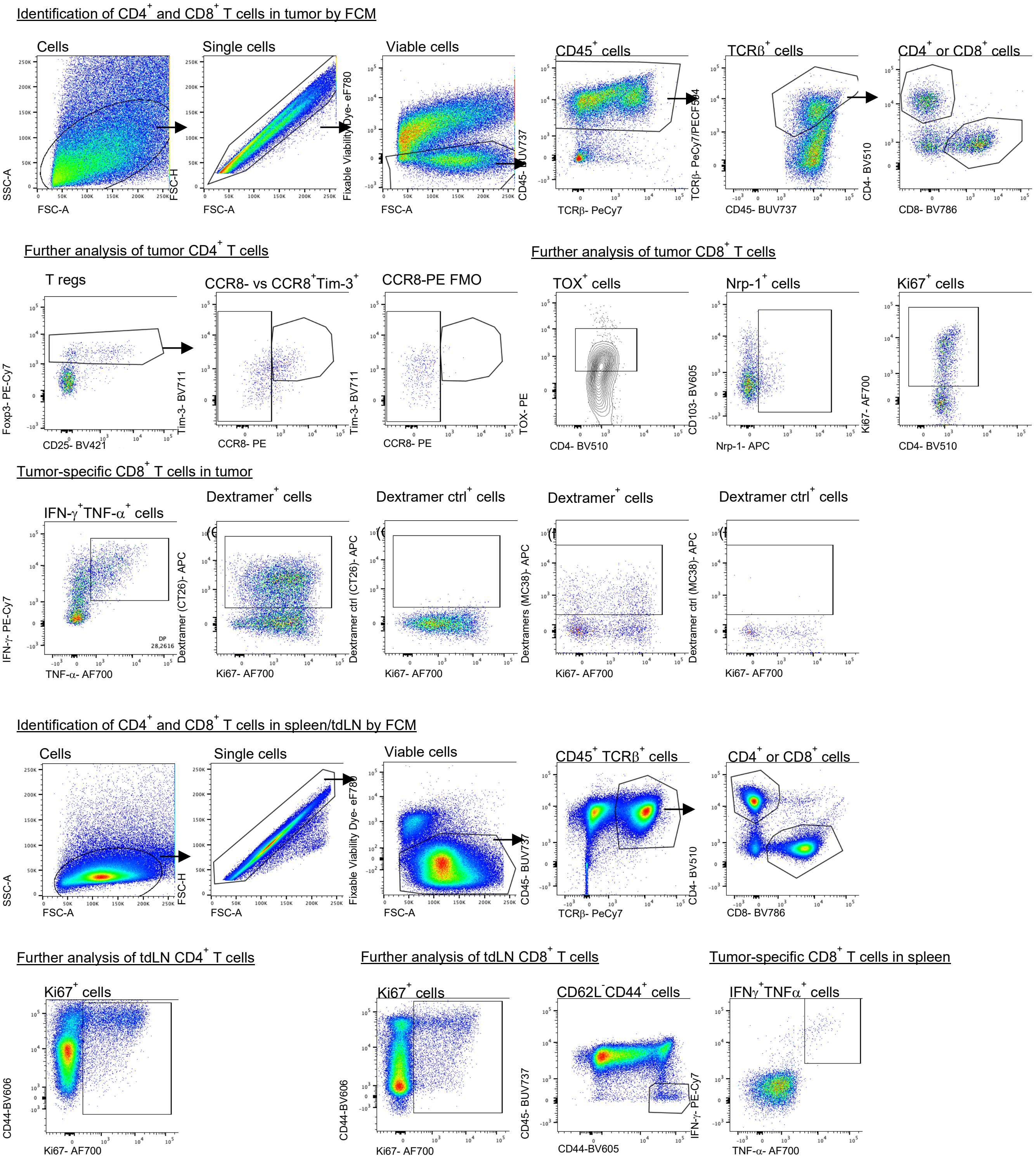

Supplementary figure 1: Gating strategy for T cells in ex vivo isolated murine tumors

Flow cytometric gating strategy to define different T cell populations in tumors and lymph nodes (LNs). In both tumor and LNs, lymphocytes were identified based on FSC-A and SSC-A, singlets by FSC-A and FSC-H, dead cells were excluded using a viability dye, $CD45^{+}$ cells expressing TCRb identified T cells which then were divided into CD4 or CD8 expressing cells. In the tumor sample, T regs were identified in the $CD4^{+}$ T cells by gating on Foxp3 expressing cells. T regs were then divided into $CCR8^{+}$ and $CCR8^{-}$ cells. Proliferating T cells were identified using Ki67. Levels of the exhaustion markers TOX and Neuropilin-1 (Nrp-1) were assessed for tumor $CD8^{+}$ T cells. Tumor-specific $CD8^{+}$ T cells were identified using MHC I dextramer reagents or by assessing IFN-g and TNF-a producing $CD8^{+}$ T cells after culture with tumor peptides. In LNs, $CD8^{+}$ T cells expressing CD44 but not CD62L were identified as effector memory cells.

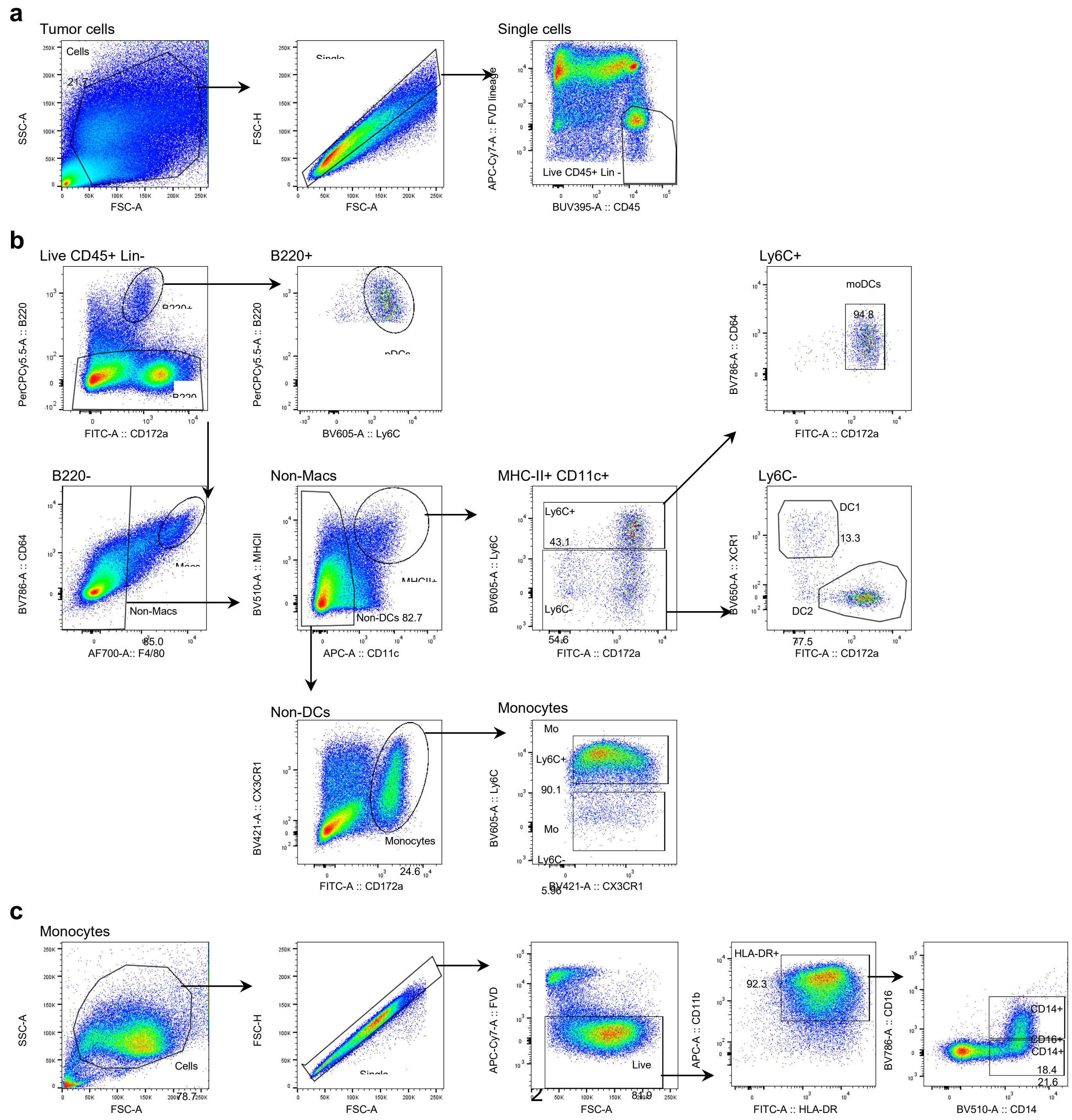

Supplementary figure 2: Gating strategy for macrophages, monocytes and dendritic cells from murine tumors at 8 days and for monocytes in human PBMC

**a,** Single cell suspensions derived from harvested tumors at D8 post treatment were evaluated by FCM. Prior to concatenation of CD45+ tumor infiltrating cells, cells were gated based on forward and side scatters followed by doublet elimination. Dead cells and lineage positive cells (TCRβ, CD19, NK1,1, CD49b, SiglecF and Ly6G) were excluded and only CD45+ live lineage negative cells were selected for concatenation and further analysis. **b,** Different myeloid cells can be identified by surface expression markers and manual gating of tumor infiltrating CD45+ live lineage negative cells. pDCs were gated based on B220 and dim Ly6C+ expression. All B220- myeloid cells were then gated based on their expression of CD64 and F4/80, discriminating between Macrophages (Macs, CD64+ F4/80+) and non-macrophages (non-Macs, F4/80-). Within non-macrophage populations, conventional DCs and monocyte-derived DCs (moDCs) were gated within MHC-II+ and CD11c+ cells and discriminated based on Ly6C expression. MoDCs were gated within MHC-II+ CD11c+ and express Ly6C+ CD64+ CD172a+. Conventional dendritic cells type 1 (DC1) and type 2 (DC2) are Ly6C- CD172a- XCR1+ or Ly6C- CD172a+ XCR1-, respectively. Monocytes were gated outside of MHC-II+ CD11c+ as belonging to the non-DCs gate. To restrict the population further, only cells expressing CD172a+ were considered Monocytes that then are separated in Ly6C+ monocytes (Mo Ly6C+) and Ly6C- monocytes (Mo Ly6C-). **c,** Human PBMCs were harvested from leucocyte concentrates and CD14+ monocytes isolated by positive selection prior to culture with anti-TNFR2 antibody or isotype control for 20h. FCM gating strategy is depicted. After exclusion of doublets and dead cells, HLA-DR+ CD11b+ were separated in CD14+ and CD14+ CD16+ monocytes for further analysis.

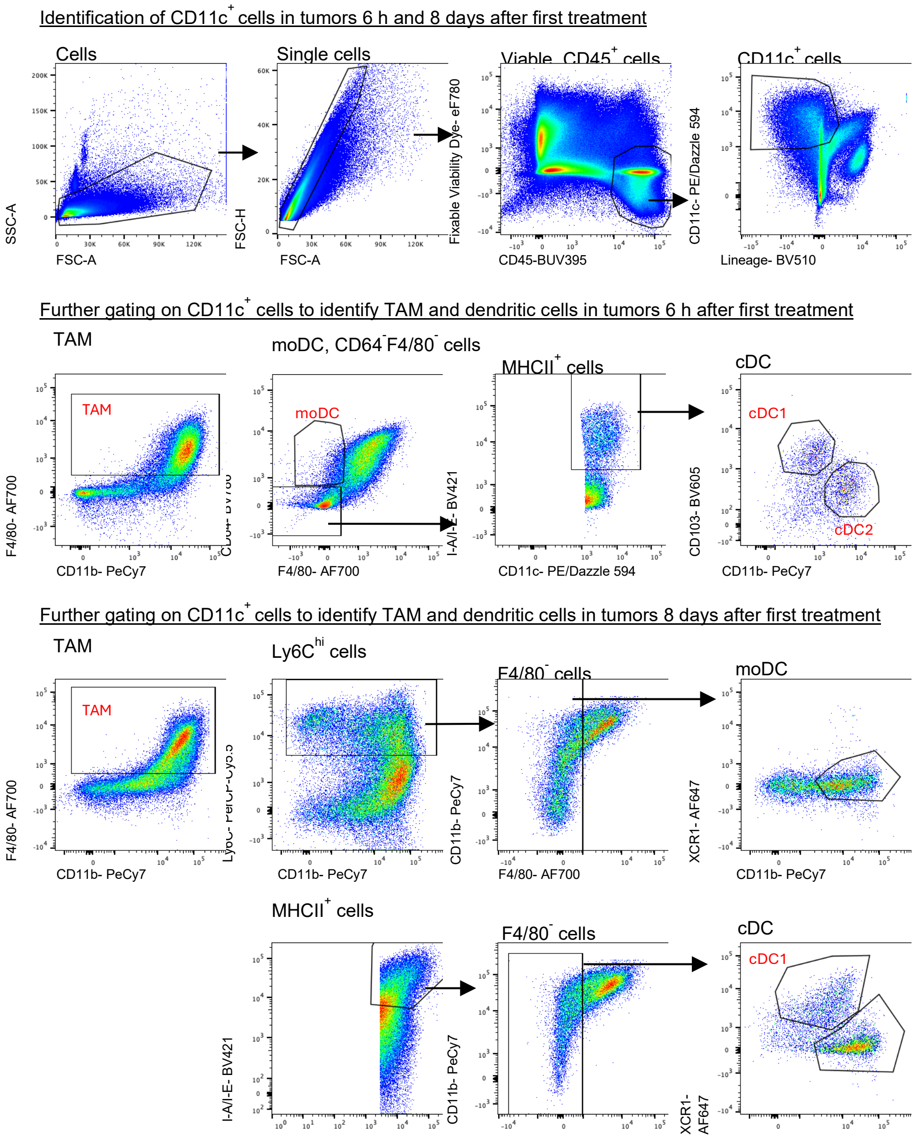


Supplementary figure 3: Gating strategy for macrophages and dendritic cells from murine tumors at 6 h

Gating strategy to identify tumor associated macrophages (TAMs), monocyte-derived dendritic cells (Mo-DCs), conventional DC1s (cDC1s) and cDC2s in tumors and tumor draining lymph nodes (tdLNs). Organs were harvested 6 h after first treatment and were gated on cells based on FSC-A and SSC-A, singlets by FSC-A and FSC-H, dead cells were excluded using a viability dye and then gated on $CD45^+$ cells. From $CD45^+$ cells, $CD11c^+$lineage (TCRVb3, NK1.1, CD19 and siglecF)$^-$ cells were identified. TAMs were identified within the $CD11c^+$ population gating on $F4/80^+CD11b^+$ cells. MoDCs were identified from the $CD11c^+$ cells as $F4/80^-$ and $CD64^+$ and cDCs as $F4/80^-CD64^-$ and $MHCII^+$. cDCs were then further divided into cDC1 and cDC2 based on CD103 and CD11b expression (cDC1: $CD103^+CD11b^-$ and cDC2: $CD103^-CD11b^+$).

Flow cytometry antibody panels (FCM panels)

1. Binding in vitro activated CD4⁺ T cells

*Human*

| Color | Antigen | Clone | Company | Cat. # |
|---|---|---|---|---|
| eFlour780 | eBioscience™ Fixable Viability Dye | | ThermoFisher | 65-0865-18 |
| BV510 | CD4 | L200 | BD | 563094 |
| BV421 | CD25 | M-A251 | BioLegend | 356114 |
| APC | AffiniPure™ F(ab')$_2$ Fragment Donkey Anti-Human IgG (H+L) | Polyclonal | Jackson Immunoresearch | 709-136-149 |

*Mouse*

| | | | | |
|---|---|---|---|---|
| eFlour780 | eBioscience™ Fixable Viability Dye | | ThermoFisher | 65-0865-18 |
| BV510 | CD4 | RM4-5 | BD | 563106 |
| BV421 | CD25 | PC61 | BioLegend | 102034 |
| APC | AffiniPure™ F(ab')$_2$ Fragment Donkey Anti-Mouse IgG (H+L) | Polyclonal | Jackson Immunoresearch | 715-136-151 |

2. T cell activation assay

| Color | Antigen | Clone | Company | Cat. # |
|---|---|---|---|---|
| eFlour780 | eBioscience™ Fixable Viability Dye | | ThermoFisher | 65-0865-18 |
| BV510 | CD4 | L200 | BD | 563094 |
| Pe-Cy7 | CD8 | RPA-T8 | BD | 557746 |
| BV421 | CD25 | M-A251 | BioLegend | 356114 |
| CFSE | | | Invitrogen | C34554 |

3. TAM-T cell co-culture

*Human*

| Color | Antigen | Clone | Company | Cat. # |
|---|---|---|---|---|
| eFlour780 | eBioscience™ Fixable Viability Dye | N/A | ThermoFisher | 65-0865-18 |
| BV421 | CD25 | M-A251 | BD | 562442 |
| CFSE | | | Invitrogen | C34554 |

*Mouse*

| Color | Antigen | Clone | Company | Cat. # |
|---|---|---|---|---|
| eFlour780 | eBioscience™ Fixable Viability Dye | | ThermoFisher | 65-0865-18 |
| BV421 | CD25 | PC61 | BioLegend | 102034 |
| CFSE | | | Invitrogen | C34554 |

4. TNFR2 chimeric mice

*CD4 and CD8 reconstitution*

| Color | Antigen | Clone | Company | Cat. # |
|---|---|---|---|---|
| FITC | CD45 | 30-F11 | BD | 553080 |
| APC | TCRb | H57-597 | aviva | OATA00096 |
| BV510 | CD4 | RM4-5 | BD | 563106 |
| PeCy7 | CD8 | 53-6.7 | BD | 552877 |
| eFlour780 | eBioscience™ Fixable Viability Dye | | ThermoFisher | 65-0865-18 |

*Human and mouse TNFR2 expression*

| | | | | |
|---|---|---|---|---|
| BV421 | CD45 | 30-F11 | BioLegend | 103133 |
| AF488 | TCRb | H57-597 | BioLegend | 109215 |
| BV605 | Ly6C | AL-21 | BD | 563011 |
| BV711 | CD11b | M1/70 | BioLegend | 101242 |
| APC | Streptavidin | | Miltenyi | 130-106-791 |
| Biotin | in-house TNFR2 (BI-1808 (human) or 3-F10 (mouse) | | | |
| eFlour780 | eBioscience™ Fixable Viability Dye | | ThermoFisher | 65-0865-18 |

5. Confirmation of CD8 depletion

| Color | Antigen | Clone | Company | Cat. # |
|---|---|---|---|---|
| PE-Cy7 | CD45.2 | 104 | Biolegend | 109830 |
| Pacific Blue | CD8a | 53-6.7 | Biolegend | 100725 |
| APC-Cy7 | CD4 | GK1.5 | Biolegend | 100414 |
| PerCP-Cyanine5. | CD19 | 1D3 | Biolegend | 152403 |

6. Sort scRNAseq

| Color | Antigen | Clone | Company | Cat. # |
|---|---|---|---|---|
| eFlour780 | eBioscience™ Fixable Viability Dye | | ThermoFisher | 65-0865-18 |
| BV421 | CD45 | 30-F11 | BioLegend | 103133 |

7. Identification of responding animals (scRNAseq)

| Color | Antigen | Clone | Company | Cat. # |
|---|---|---|---|---|
| eFlour780 | eBioscience™ Fixable Viability Dye | | ThermoFisher | 65-0865-18 |
| BUV395 | CD62L | MEL-14 | BD | 740218 |
| BUV737 | CD45.2 | 104 | BD | 612779 |
| BV421 | CX3CR1 | SA011F11 | BioLegend | 149023 |
| BV510 | CD4 | RM4-5 | BD | 563106 |
| BV605 | CD44 | IM7 | BD | 563058 |
| BV786 | CD8a | 53-6.7 | BD | 563332 |
| AF488 | TCRb | H57-597 | BioLegend | 109215 |
| APC | MHC class I Dextramers | | Immudex | JG3294-APC |

8. TNFR2 immune cell expression- mouse tumor and blood

*T cells*

| Color | Antigen | Clone | Company | Cat. # |
|---|---|---|---|---|
| PE | TNFR2 | 55R-286/TR75-89 | Biolegend/BioLegend | 113004/113406 |
| AF488 | FoxP3 | FJK-16s | Invitrogen / ThermoFisher Scientific | 53-5773-82 |
| APC | PD-1 | 29F.1A12 | Biolegend | 135210 |
| PE-Cy7 | CD45.2 | 104 | Biolegend | 109830 |
| APC-Cy7 | CD4 | GK1.5 | Biolegend | 100414 |
| Pacific Blue | CD8a | 53-6.7 | Biolegend | 100725 |

*B cells and NK cells*

| Color | Antigen | Clone | Company | Cat. # |
|---|---|---|---|---|
| PE | TNFR2 | 55R-286/TR75-89 | Biolegend/BioLegend | 113004/113406 |
| PerCP-Cyanine5.5 | B220 | RA3-6B2 | Biolegend | 103236 |
| APC | NKp46 | 29A1.4 | Biolegend | 137608 |
| PE-Cy7 | CD45.2 | 104 | Biolegend | 109830 |
| eFlour 506 | Viability | - | Invitrogen / ThermoFisher Scientific | 65-0866-14 |

*Myeloid cells*

| Color | Antigen | Clone | Company | Cat. # |
|---|---|---|---|---|
| PE | TNFR2 | 55R-286/TR75-89 | Biolegend/BioLegend | 113004/113406 |
| FITC | CD11c | N418 | Biolegend | 117306 |
| PerCP-Cy5.5 | Ly6C | HK1.4 | Biolegend | 128012 |
| APC | F4/80 | BM8 | eBioscience/ThermoFisher Scientific | 17-4801-82 |
| PE-Cy7 | CD45.2 | 104 | Biolegend | 109830 |
| APC-Cy7 | Ly6G | 1A8 | Biolegend | 127624 |
| eFlour 506 | Viability | - | Invitrogen / ThermoFisher Scientific | 65-0866-14 |
| Pacific Blue | CD11b | M1/70 | Biolegend | 101224 |

9. Immune profiling Panel 1 (tumor and lymph node)

| Color | Antigen | Clone | Company | Cat. # |
|---|---|---|---|---|
| eFlour780 | eBioscience™ Fixable Viability Dye | | ThermoFisher | 65-0865-18 |
| BUV395 | CD62L | MEL-14 | BD | 740218 |
| BUV737 | CD45.2 | 104 | BD | 612779 |
| BV421 | CD25 | 7D4 | BD | 564571 |
| BV510 | CD4 | RM4-5 | BD | 563106 |
| BV605 | CD103 | 2E7 | BioLegend | 121433 |
| BV711 | Tim-3 | RMT3-23 | BioLegend | 119727 |
| BV786 | CD8a | 53-6.7 | BD | 563332 |
| AF488 | TCF | C63D9 | Cell Signaling Technology | 6444S |
| BB700 | PD-1 | RMP1-30 | BD | 748242 |
| PECF594 | TCRb | H57-597 | BD | 562841 |
| PeCy7 | Foxp3 | FJK-16s | ThermoFisher | 25-5773-82 |
| APC | NRP-1 | 3E12 | BioLegend | 145206 |
| AF700 | ki-67 | SolA15 | ThermoFisher | 56-5698-82 |
| PE | CCR8 | SA214G2 | BioLegend | 150312 |

#### 10. Immune profiling Panel 2

*Tumor*

| Color | Antigen | Clone | Company | Cat. # |
|---|---|---|---|---|
| eFlour780 | eBioscience™ Fixable Viability Dye | | ThermoFisher | 65-0865-18 |
| BUV395 | CD62L | MEL-14 | BD | 740218 |
| BUV737 | CD45.2 | 104 | BD | 612779 |
| BV421 | GzmB | GB11 | BD | 563389 |
| BV510 | CD4 | RM4-5 | BD | 563106 |
| BV605 | CD103 | 2E7 | BioLegend | 121433 |
| BV711 | Tim-3 | RMT3-23 | BioLegend | 119727 |
| BV786 | CD8a | 53-6.7 | BD | 563332 |
| AF488 | TCF | C63D9 | Cell Signaling Technology | 6444S |
| BB700 | PD-1 | RMP1-30 | BD | 748242 |
| PE | TOX | TXRX10 | ThermoFisher | 12-6502-80 |
| PECF594 | CD39 | Y23-1185 | BD | 567268 |
| PeCy7 | TCRb | H57-597 | BioLegend | 109222 |
| APC | MHC class I Dextramers (CT26 or MC38) | | Immudex | |
| AF700 | ki-67 | SolA15 | ThermoFisher | 56-5698-82 |

*Lymph node*

| Color | Antigen | Clone | Company | Cat. # |
|---|---|---|---|---|
| eFlour780 | eBioscience™ Fixable Viability Dye | | ThermoFisher | 65-0865-18 |
| BUV395 | CD62L | MEL-14 | BD | 740218 |
| BUV737 | CD45.2 | 104 | BD | 612779 |
| BV421 | GzmB | GB11 | BD | 563389 |
| BV510 | CD4 | RM4-5 | BD | 563106 |
| BV605 | CD44 | IM7 | BD | 563058 |
| BV711 | CX3CR1 | SA011F11 | BioLegend | 149031 |
| BV786 | CD8a | 53-6.7 | BD | 563332 |
| AF488 | TCF | C63D9 | Cell Signaling Technology | 6444S |
| BB700 | PD-1 | RMP1-30 | BD | 748242 |
| PE | TOX | TXRX10 | ThermoFisher | 12-6502-80 |
| PECF594 | CD39 | Y23-1185 | BD | 567268 |
| PeCy7 | TCRb | H57-597 | BioLegend | 109222 |
| APC | MHC class I Dextramers (CT26 or MC38) | | Immudex | |
| AF700 | ki-67 | SolA15 | ThermoFisher | 56-5698-82 |

#### 11. CD86 expression on TAM, moDC, cDC1 and cDC2, day 8

| Color | Antigen | Clone | Company | Cat. # |
|---|---|---|---|---|
| BUV395 | CD45 | 30-F11 | BD | 564279 |
| BUV737 | CD86 | GL1 | BD | 741737 |
| BV421 | CX3CR1 | Z8-50 | BD | 567531 |
| BV510 | MHC-II | M5/114.15.2 | BioLegend | 107635 |
| BV605 | Ly6c | HK1.4 | BioLegend | 128036 |
| BV650 | XCR1 | ZET | BioLegend | 148220 |
| BV786 | CD64 | X54-5/7.1 | BD | 569507 |

| | | | | |
|---|---|---|---|---|
| AF488 | CD172a | P84 | BioLegend | 144024 |
| PerCP-Cy5.5 | B220 | RA3-6B2 | BD | 557271 |
| PE | CD206 | Y17-505 | BD | 568273 |
| PE-Cy7 | CD11b | M1/70 | BD | 552850 |
| APC | CD11c | N418 | BioLegend | 117310 |
| AF700 | F4/80 | BM8 | BioLegend | 123130 |
| APC-Cy7 | CD49b | DX5 | BioLegend | 108920 |
| APC-Cy7 | SiglecF | E50-2440 | BD | 565527 |
| APC-Cy7 | Ly6G | 1A8 | BD | 560600 |
| APC-Cy7 | TCRb | H57-597 | BioLegend | 109220 |
| APC-Cy7 | CD19 | 1D3 | BioLegend | 152412 |
| APC-Cy7 | NK1.1 | PK136 | BD | 560618 |
| eFluor780 | eBioscience Fixable Viability Dye | | ThermoFisher | 65-0865-18 |

#### 12. CD86 expression on TAM, moDC, cDC1 and cDC2, 6 h

| Color | Antigen | Clone | Company | Cat. # |
|---|---|---|---|---|
| BV421 | I-A/I-E | M5/114.15.2 | BD | 562564 |
| BV510 | NK1.1 (Lineage) | PK136 | BD | 563096 |
| BV510 | CD19 (Lineage) | 1D3 | BD | 562956 |
| BV510 | TCRVb3 (Lineage) | KJ25 | BD | 743413 |
| BV510 | Siglec F (Lineage) | E50-2440 | BD | 740158 |
| BV605 | CD103 | 2E7 | BioLegend | 121433 |
| BV786 | CD64 | X54-5/7.1 | BD | 741024 |
| BUV395 | CD45 | 30-F11 | BD | 564279 |
| BUV737 | CD86 | GL1 | BD | 741737 |
| AF700 | F4/80 | BM8 | BioLegend | 123130 |
| eFlour780 | eBioscience™ Fixable Viability Dye | | ThermoFisher | 65-0865-18 |
| PE/Dazzle™ 594 | CD11c | N418 | BioLegend | 117348 |
| PECy7 | CD11b | M1/70 | BD | 552850 |

#### 13. Antigen-specific CD8 T cells- cultures with peptide

| Color | Antigen | Clone | Company | Cat. # |
|---|---|---|---|---|
| eFlour780 | eBioscience™ Fixable Viability Dye | | ThermoFisher | 65-0865-18 |
| AF488 | TCRb | H57-597 | BioLegend | 109215 |
| BV510 | CD4 | RM4-5 | BD | 563106 |
| BV786 | CD8a | 53-6.7 | BD | 563332 |
| BV421 | CD25 | PC61 | BioLegend | 102034 |
| BUV395 | CD45 | 30-F11 | BD | 564279 |
| PeCy7 | IFN-g | XMG1.2 | BioLegend | 505826 |
| AF700 | TNF-a | MP6-XT22 | BD | 558000 |

#### 14. TNFR2 immune cell expression- human blood

*T cell populations*

| Fluorochromes | Antigen | Clone | Company | Cat. # |
|---|---|---|---|---|
| BUV395 | CD45 | HI30 | BD | 56792 |
| BV510 | CD4 | L200 | BD | 563094 |
| BV711 | CCR7 | 150503 | BD | 566602 |
| BV785 | CD8 | RPA-T4 | BioLegend | 301046 |
| AF488 | CD3 | UCHT-1 | BD | 557694 |
| PE | CD25 | BC96 | BD | 567214 |
| PE-Vio770 | CD45RO | UCHL1 | Miltenyi | 130-113-551 |
| Biotin | TNFR2 | BI-1808 | in-house | |
| APC | Streptavidin | | Miltenyi | 130-106-791 |
| APC-R700 | CD127 | HIL-7R-M21 | BD | 565185 |
| eFlour780 | eBioscience™ Fixable Viability Dye | | ThermoFisher | 65-0865-18 |

*NK cells, B cells, granulocytes and monocytes*

| Fluorochromes | Antigen | Clone | Company | Cat. # |
|---|---|---|---|---|
| BUV395 | CD45 | HI30 | BD | 56792 |
| BUV737 | CD19 | SJ25C1 | BD | 612757 |
| BV510 | CD14 | 63D3 | BioLegend | 367124 |
| BV711 | CD33 | WM53 | BD | 563171 |
| BV786, | CD16 | 3G8 | BD | 563690 |
| AF488 | CD3 | UCHT-1 | BD | 557694 |
| PE | CD56 | MY31 | BD | 556647 |
| PECy7 | CD123 | 9F5 | BD | 551065 |
| Biotin | TNFR2 | BI-1808 | in-house | |
| APC | Streptavidin | | Miltenyi | 130-106-791 |
| AF700 | CD66 | | | |
| eFlour780 | eBioscience™ Fixable Viability Dye | | ThermoFisher | 65-0865-18 |

#### 15. Human monocyte cultures

| Color | Antigen | Clone | Company | Cat. # |
|---|---|---|---|---|
| BV510 | CD14 | 63D3 | BioLegend | 367124 |
| BV786 | CD16 | 3G8 | BD | 563690 |
| BB515 | HLA-DR | G46-6 | BD | 564516 |
| PE-Cy7 | CD86 | FUN-1 | BD | 561128 |
| APC | CD11b | M1/70 | BD | 553312 |
| eFluor780 | eBioscience Fixable Viability Dye | | ThermoFisher | 65-0865-18 |
